# Analysis of independent cohorts of outbred CFW mice reveals novel loci for behavioral and physiological traits and identifies factors determining reproducibility

**DOI:** 10.1101/2021.02.05.429998

**Authors:** Jennifer Zou, Shyam Gopalakrishnan, Clarissa C. Parker, Jerome Nicod, Richard Mott, Na Cai, Arimantas Lionikas, Robert W Davies, Abraham A. Palmer, Jonathan Flint

## Abstract

Combining samples for genetic association is standard practice in human genetic analysis of complex traits, but is rarely undertaken in rodent genetics. Here, using 23 phenotypes and genotypes from two independent laboratories, we obtained a sample size of 3,076 commercially available outbred mice and identified 70 loci, more than double the number of loci identified in the component studies. Fine-mapping in the combined sample reduced the number of likely causal variants, with a median reduction in set size of 51%, and indicated novel gene associations, including Pnpo, Ttll6 and GM11545 with bone mineral density, and Psmb9 with weight. However replication at a nominal threshold of 0.05 between the two component studies was surprisingly low, with less than a third of loci identified in one study replicated in the second. In addition to overestimates in the effect size in the discovery sample (Winner’s Curse), we also found that heterogeneity between studies explained the poor replication, but the contribution of these two factors varied among traits. Available methods to control Winner’s Curse were contingent on the power of the discovery sample, and depending on the method used, both overestimated and underestimated the true effect. Leveraging these observations we integrated information about replication rates, confounding, and Winner’s Curse corrected estimates of power to assign variants to one of four confidence levels. Our approach addresses concerns about reproducibility, and demonstrates how to obtain robust results from mapping complex traits in any genome-wide association study.

## Introduction

Combining samples, through meta or mega-analysis, has become routine in human genome-wide association studies (GWAS) of complex traits as a way to augment power by increasing sample size and to ensure robustness of results by replicating findings. Genetic mapping of complex traits in rodents has favored the use of linkage analysis in crosses between inbred strains, with many different inbred strain combinations being employed. In addition, few studies examine the same phenotypes. Therefore, rodent studies have not lent themselves to meta- or mega-analysis (Wuschke et al. 2007; Schmidt et al. 2008). However, the more recent transition to genetic association using outbred mice (Parker and Palmer 2011; Flint and Eskin 2012; Chesler 2014) or panels of inbred animals (Ghazalpour et al. 2012) provides opportunities for deploying meta- (Kang et al. 2014) and mega-analysis (Chitre et al. 2020; Zhou et al. 2020) to increase power and test reproducibility.

In this paper we combine results from two independent laboratories, one at the University of Oxford (OX) in the United Kingdom (Nicod et al. 2016) and one at the University of Chicago (UC) in the United States of America (Parker et al. 2016). Both experiments sampled from the same population of commercially available outbred mice (Crl:CFW(SW)-US_P08, hereafter CFW), but differed in genotyping platforms, and in some of the phenotyping assays. The datasets provided an opportunity to examine several questions that have not been fully addressed in mouse GWAS.

While a consensus has arisen in human GWAS that a 5×10^−8^ threshold together with replication in an independent sample is sufficient to declare locus discovery (Pe’er et al. 2008; Visscher et al. 2012), the same is not true for mouse studies. Mouse populations used for GWAS differ substantially, in linkage disequilibrium (LD) structure, allele frequencies and the extent of relatedness between subjects, making it inappropriate to codify a single significance threshold (Flint and Eskin 2012). Furthermore, the selection of a genome-wide significance threshold introduces a bias known as Winner’s Curse, in which loci passing the genome-wide significance threshold tend to have inflated effect sizes (Zhong and Prentice 2008; Sun et al. 2011b; Xiao and Boehnke 2011). Winner’s Curse contributes substantially to replication of GWAS loci in follow-up studies (Palmer and Pe’er 2017). There have also been concerns about the impact of laboratory differences on the measurement of behavior (Crabbe et al. 1999) and the potentially large impact of confounding by subtle laboratory effects (Valdar et al. 2006; Zhou et al. 2020). These differences between studies can contribute to study-specific heterogeneity, which can lead to spurious associations when combining studies. These issues question the generalizability of genetic analyses.

Here we performed a mega-analysis between the OX and UC cohorts and identified 70 independent loci, of which 41 were not found in component studies. Novel loci can result from the increase in power from combining data in a mega-analysis, or they can result from study-specific heterogeneity between the component studies. To investigate the robustness of the novel associations in the mega-analysis, we integrate assessments of heritability, genetic correlation, locus-specific replication, estimates of study-specific heterogeneity (Zou et al. 2020), and Winner’s Curse corrected estimates of power (Zhong and Prentice 2008) to categorize loci into one of four confidence levels. We then performed fine-mapping analysis and annotation of nonsynonymous variants to identify putative genes for a number of phenotypes.

## Results

We provide an overview of our methodology in Figure 1, which shows the numbers of phenotypes, QTLs, and genes that we identified at each stage of the analysis.

**Figure 1:**
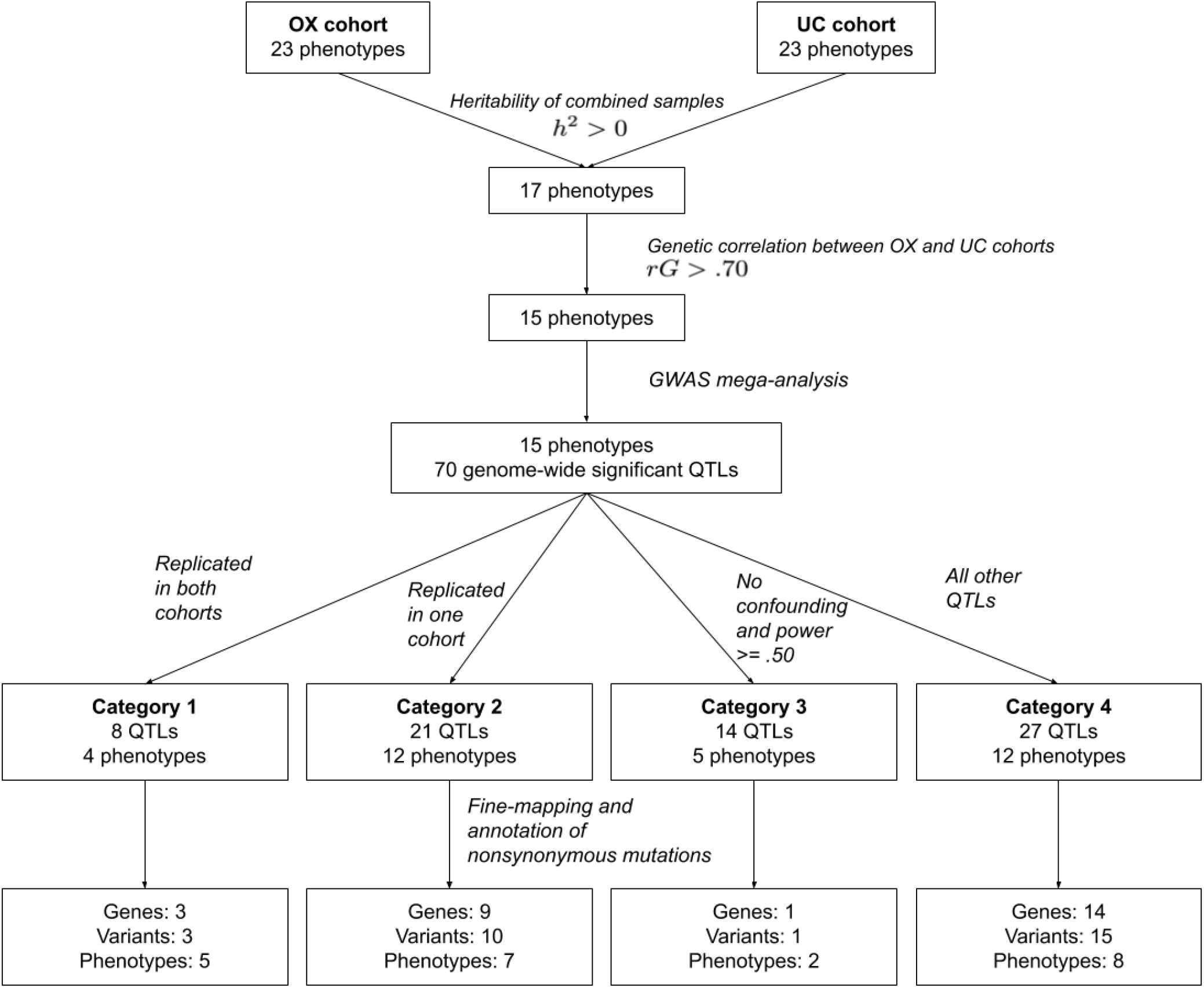
Method overview. We combined two cohorts of CFW mice for 23 shared phenotypes. We identified 17 phenotypes with heritability significantly greater than zero. 15 of these phenotypes had genetic correlations greater than .70. We performed GWAS meta-analysis using these 15 phenotypes and obtained 70 genome-wide significant QTLs. We then integrated replication data, estimates of confounding, and winner’s curse corrected estimates of power to categorize the 70 variants into one of four categories. Finally, we performed fine-mapping and annotation of nonsynonymous mutations to identify candidate genes.

### Combining phenotypes and genotypes

We identified 23 phenotypes that were measured in both studies (Supplemental Table 1 and Supplemental Data 1). We observed a high correlation between related phenotypes, such as “locomotor activity initial” and “locomotor activity total” (Supplemental Figure 1). To obtain a common set of genotypes, we converted sequence data to genotypes using a single pipeline and obtained a common set of 3,152,108 SNPs for mapping with MAF > 0.001. Quality control data for genotypes are shown in Supplemental Figure 2. We also genotyped a subset of these mice on the megaMUGA array (Collaborative Cross 2012), and observed 98.71% concordance for the OX cohort and 97.14% concordance for the UC cohort. We also compared the overlapping SNPs to our prior publications and found 99.12% concordance for OX and 91.87% concordance for UC. After filtering the imputed genotypes for pairwise r^2^ (>0.999), 97,452 SNPs were retained for subsequent mapping (Supplemental Data 2). For all phenotypes, residuals were obtained by regressing out the relevant covariates in a linear model. These residuals were quantile normalized within each cohort and then combined for the mega analysis.

### Genetic Correlations

To determine the degree of concordance between the OX and UC component studies, we computed genetic correlations using bivariate genome-based restricted maximum likelihood estimates, implemented in the GCTA software package (Yang et al. 2011) (Figure 2). Genetic correlations were obtained for 17 of the 23 phenotypes with non-zero heritabilities (Supplemental Table 2) using samples from both studies. For these analyses, we used a set of pruned variants with INFO>0.90. As a control, we performed the same analyses using two cohorts that were obtained by randomly splitting the OX cohort in half (termed the OX1 and OX2). Theoretically, there should be little confounding between OX1 and OX2, so the genetic correlation should be close to 1.

**Figure 2:**
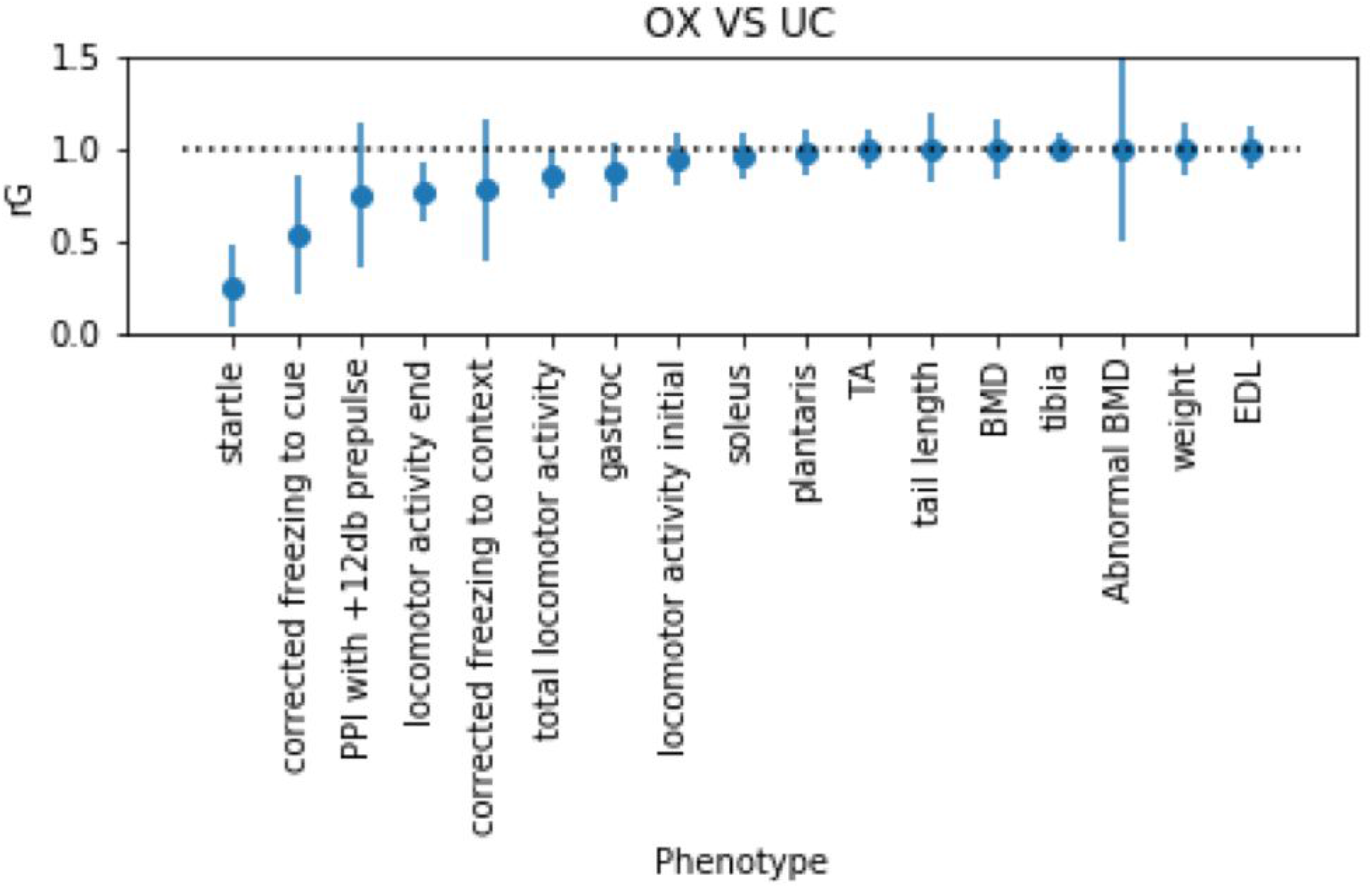
The genetic correlation between each pair of phenotypes mapped in OX and UC studies. Each dot represents the estimated genetic correlation, and the error bars show the 95% confidence intervals obtained using the standard errors. Only phenotypes with heritability significantly different from zero are shown. Phenotype names are defined in Supplemental Table 1

As expected, the genetic correlation between OX1 and OX2 was indeed close to 1 for all phenotypes (Supplemental Figure 3). The standard errors on the genetic correlations were high when the estimated heritability for the trait was low. Figure 2 shows that genetic correlations between OX and UC were greater than 0.7 for phenotypes with heritability greater than or equal to 15%, indicating that the studies have substantial shared genetic association signal. Behavioral phenotypes, such as startle and fear conditioning, tended to have lower heritabilities (Supplemental Table 2) than the physiological traits. However, this may be in part due to differences in methodology between the OX and UC studies (Methods).

### Mega-analysis

We performed a mega-analysis of 23 traits in the combined OX and UC cohorts (3,076 mice, Supplemental Data 3). We compared two different permutation-based methods (“naive” and “decorrelated”) to obtain thresholds at a significance level of 0.05 (Methods, Supplemental Figure 4). We used the more conservative decorrelated thresholds for our analysis. We identified 70 independent loci that were significant at an empirically derived −logP threshold of 5.79 (Figure 3).

**Figure 3:**
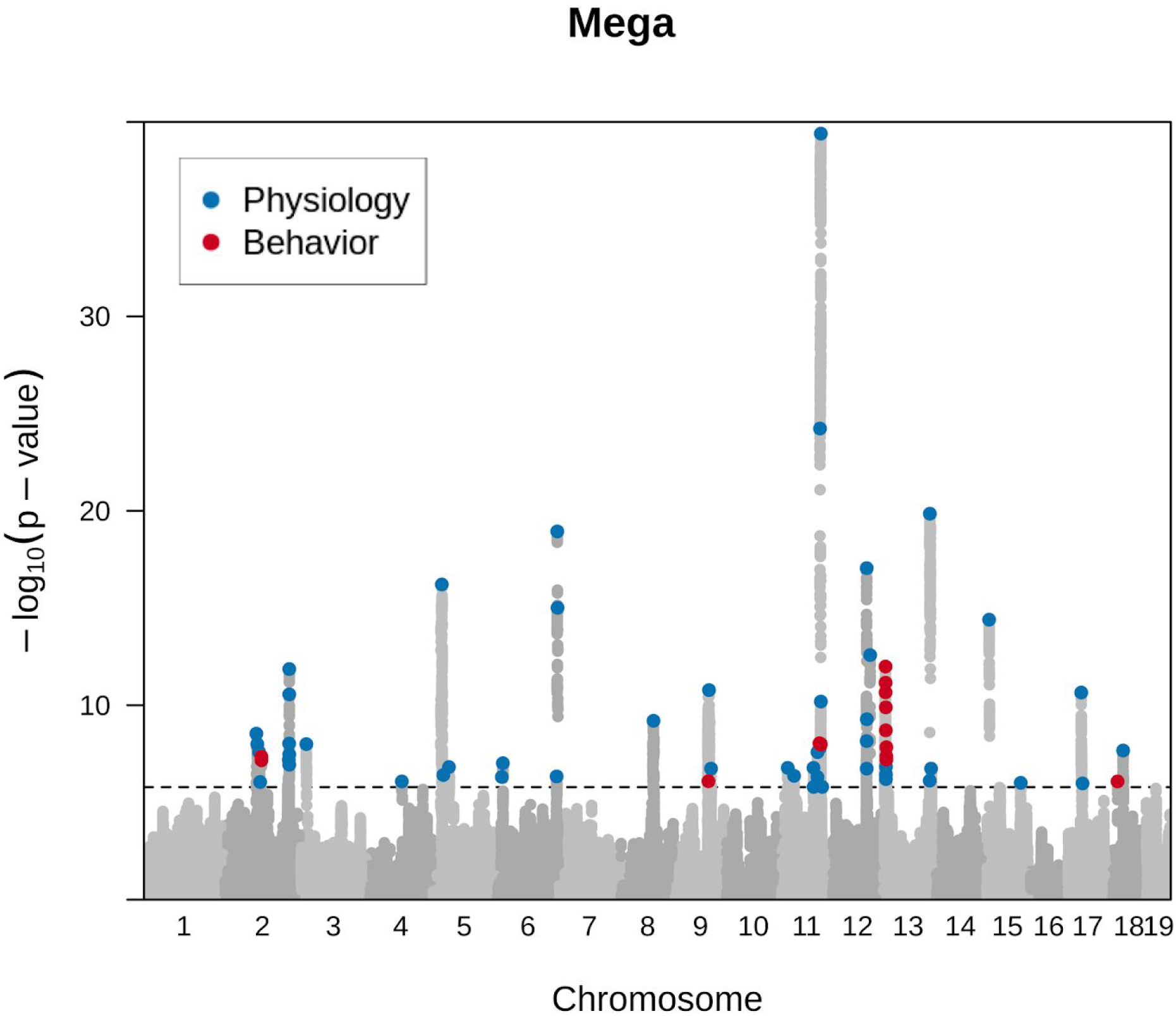
Genome-wide representation of all QTLs identified. Light grey dots show association for the measures where a QTL was detected. SNPs at each locus that exceed a permutation derived significance threshold are marked with a colored dot, where blue represents physiological and red behavioral traits. Significance thresholds were derived using a decorrelated permutation approach, described in the methods section.

The genetic architecture of the phenotypes, as illustrated in Figure 3, is consistent with high polygenicity; namely many loci each contribute a small effect to the heritability. The figure also shows that for samples of the same size, the logP values for behavior (red dots in the figure) are on average less than those for physiological traits (blue dots), revealing the relatively smaller effects at loci contributing to variation in behavior. The absolute median effect size for the behavioral loci was 0.16, and 0.18 for physiological traits. Supplemental Table 3 lists the positions of all genome-wide significant loci for every phenotype, giving their logP and effect sizes.

### Confidence intervals

We estimated the 95% confidence interval (CI) for every QTL using a simulation procedure in which, for each locus, we randomly implanted causal SNPs that matched the true QTL’s observed effect size and simulated phenotypes for all samples. A local scan of the region using the mixed model was then performed using the simulated phenotypes, and the location and p-value of the top SNP was recorded. These simulations were used to compute the empirical distribution of the change in p-value between the most highly associated SNP and the causal SNP (Δ). The 95% CI was estimated as the maximum Δ among the lowest 95% of simulations. Supplemental Table 3 contains the CIs estimated for all QTLs found in the mega-analysis. Figure 4 shows several examples of CIs computed for two QTLs. Most CIs coincide with strong LD blocks, as expected (Figure 4a). However, in some cases, the confidence intervals included variants with low correlation to the lead SNP. Thus, the confidence intervals are able to identify regions responsible for the peak signal in a less arbitrary way than simple LD thresholds (Figure 4b). Supplemental Figure 5 shows the distribution of the 95% CI widths of all QTLs, which range from 1bp-7.3Mb, with a median of 1.1Mb.

**Figure 4:**
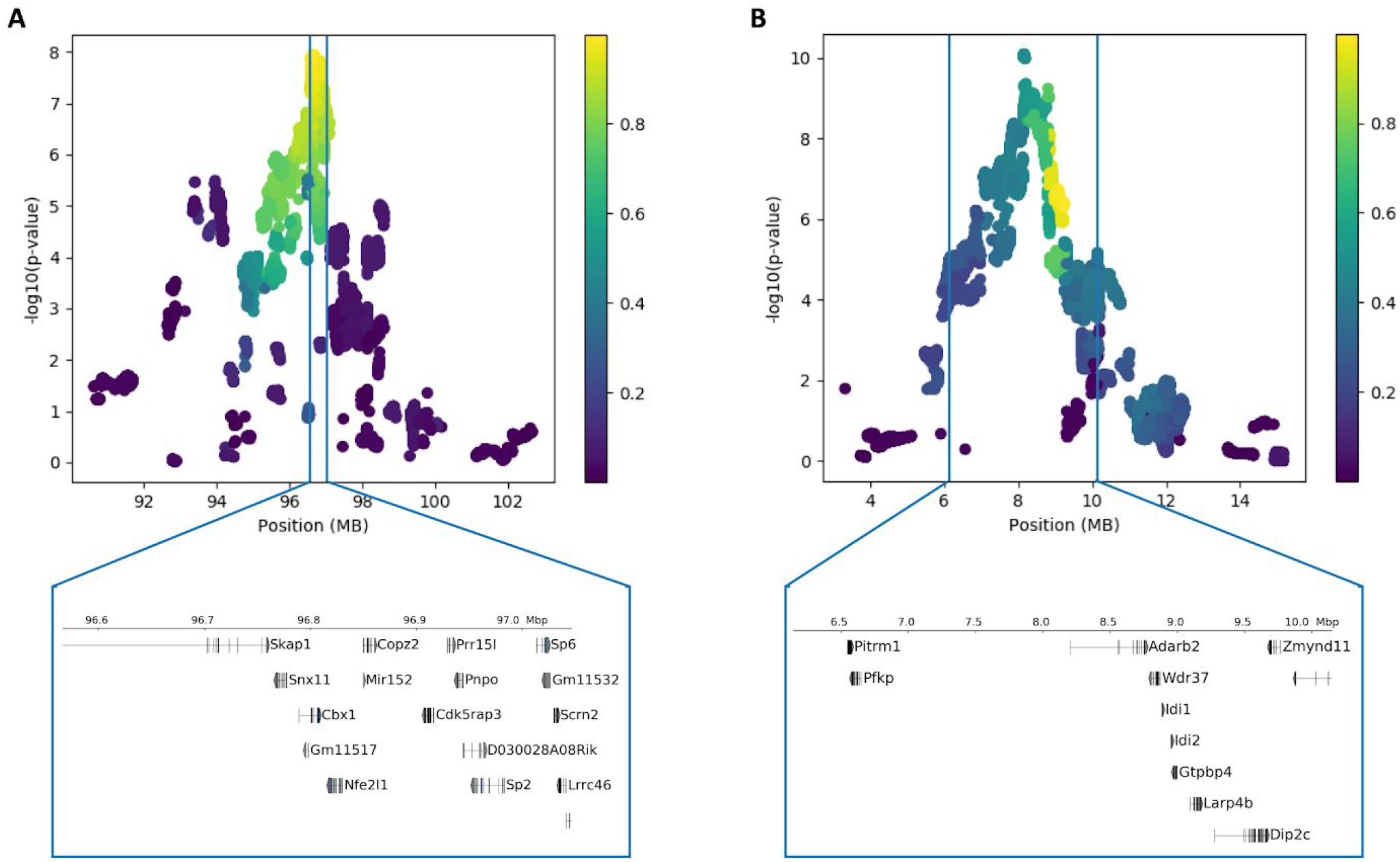
High resolution mapping of two QTLs. Each point is a variant in the locus, and the color of the point corresponds to the correlation of that variant with the lead SNP. The vertical lines correspond to the 95% confidence interval estimated for the QTL. Genes falling within the confidence intervals are shown below the plot. **A**. QTL for locomotor activity end (chr11:96653605). The confidence intervals fall along a block of variants in strong LD. **B**. QTL for initial locomotor activity (chr13:9154368). The confidence intervals include many variants with low correlation to the lead SNP..

### Fine-mapping of mega-analysis loci

In order to identify putative causal variants within the 95% CIs previously computed, we applied a fine-mapping framework called susieR (Wang et al. 2020). We used summary statistics from an LD pruned set (R2<.99) of variants and the LD between these variants as input to susieR. SusieR uses these variants to compute a number of credible sets that are designed to contain the causal variant with high probability (i.e, 95%). We obtained credible sets of SNPs for 62/70 QTLs for which the fine-mapping algorithm converged. Table 1 provides examples in which the fine-mapping was particularly effective. For example, a QTL we identified for tibia length (chr6:145481081) had over 15 thousand SNPs in the 95% CI of the QTL, and after fine-mapping, the number of putative SNPs was only 169 (representing a 99% reduction in the number of putative SNPs). The median number of variants in the credible sets was 932, and the median reduction in set size was 51% (Supplemental Figure 6, Supplemental Data 4). Across all phenotypes and loci, we identified 94,177 variants in the credible sets or in high LD (R2>.99) of variants in the credible sets. We used the variants within these credible sets as a high confidence set of variants for downstream analysis.

**Table 1:**
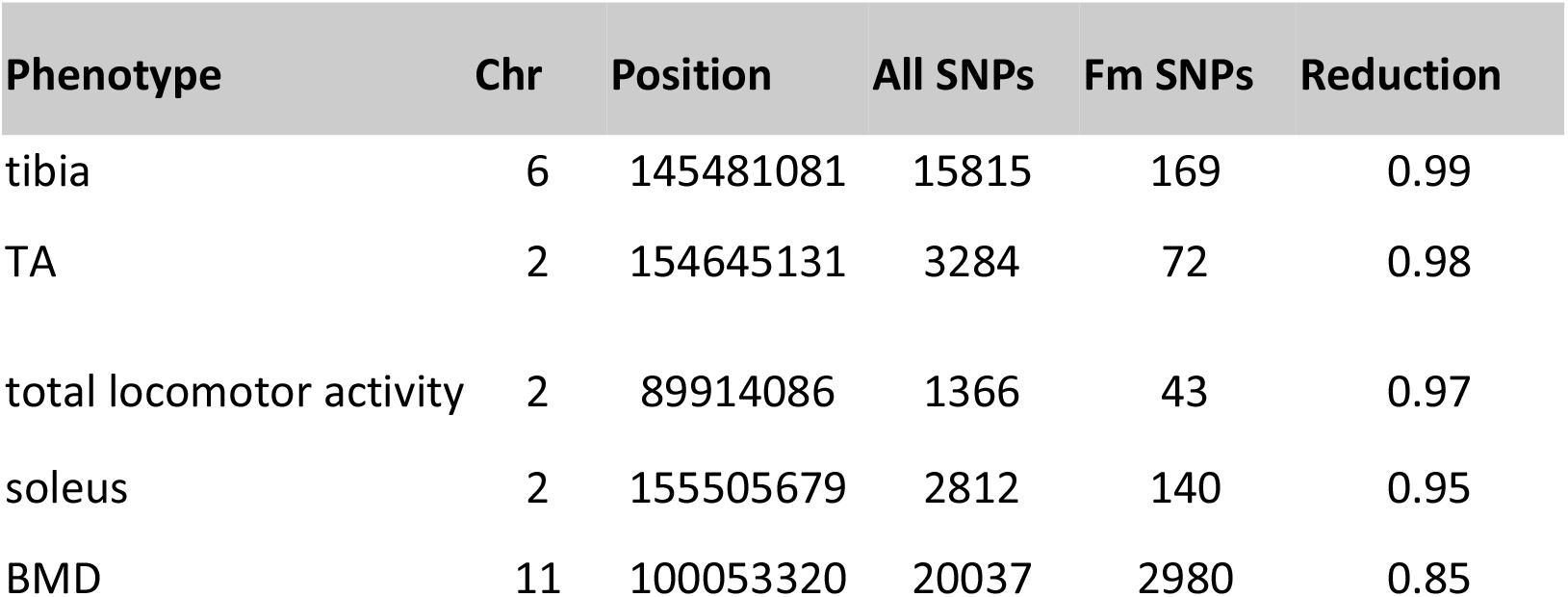
Examples of fine-mapping QTLs. “All SNPs” refers to the total number of SNPs used as input to the SusieR fine mapping method, and “Fm SNPs” corresponds to the number of variants in the credible sets. “Reduction” is the percentage of decrease relative to the total number of SNPs used as input to the fine mapping.

We generated gene-based annotations for the variants in the credible sets using the ANNOVAR software. 99% of variants were in noncoding regions of the genome. 32 variants in the credible sets were nonsynonymous mutations within 22 coding genes (compared to 4 nonsynonymous variants in UC and 16 nonsynonymous variants in OX). We interrogated the NHGRI-EBI catalog of human GWAS and the Mouse Genome Informatics database to determine if any of these 22 genes have been previously associated with relevant traits of interest Supplemental Table 4). Of the 22 coding genes identified, 11 were previously implicated in either human GWAS or KO mouse studies of similar traits (Supplemental Table 4).

### Colocalized Traits

We examined the effect of the 70 QTL significant in the mega-analysis on the other 23 traits, to determine the extent of colocalization between traits. We found that 45/70 QTLs had effects on more than one trait, after applying a Bonferroni correction (p< 5.1e-05) for multiple testing of combinations of QTLs and other phenotypes. Phenotypes often derive from the same test (such as activity measured over different lengths of time or measures obtained from different muscles). To report our results we created two phenotype categories (Behavior and Physiology). Results are shown in Supplemental Figure 7 and Supplemental Table 5. While the majority of these colocalized traits are between similar traits, such as locomotor activity at different time points, 20 of 45 QTLs had effects on at least one behavioral trait and one physiological trait. For example, a QTL for initial locomotor activity (chr11:96964818) was also associated with total locomotor activity, locomotor activity end, weight of tibialis anterior, tibia length, BMD, and abnormal BMD. Similarly, a QTL for weight (chr13:9217096) was also associated with soleus, weight of tibialis anterior, EDL, total locomotor activity, locomotor activity end, and locomotor activity initial.

### Replication

Despite the high genetic correlations between phenotypes in the OX and UC studies (Figure 2), we observed a surprisingly high rate of non-replication of QTLs, where replication was defined as a genome-wide significant association in the “discovery” dataset (OX or UC) and a p-value of 0.05 or lower in the other (“replication”) dataset. After mapping all traits in either just the OX or just the UC sample and applying empirically derived thresholds (-logp 5.68 for OX, 5.43 for UC), we identified 32 QTLs in the OX data set and 22 in the UC data set. 6 of 32 QTL discovered in OX replicated in the UC (19%); of the 22 QTL identified in UC, 7 replicated in OX (32%). There were also instances where the mega-analysis failed to confirm findings in a component study (12 of 22 loci from UC and 4 of 32 loci from OX, Supplemental Figure 8). 41 of 70 loci were significant in the mega-analysis but not significant in the component studies (Table 2).

**Table 2:**
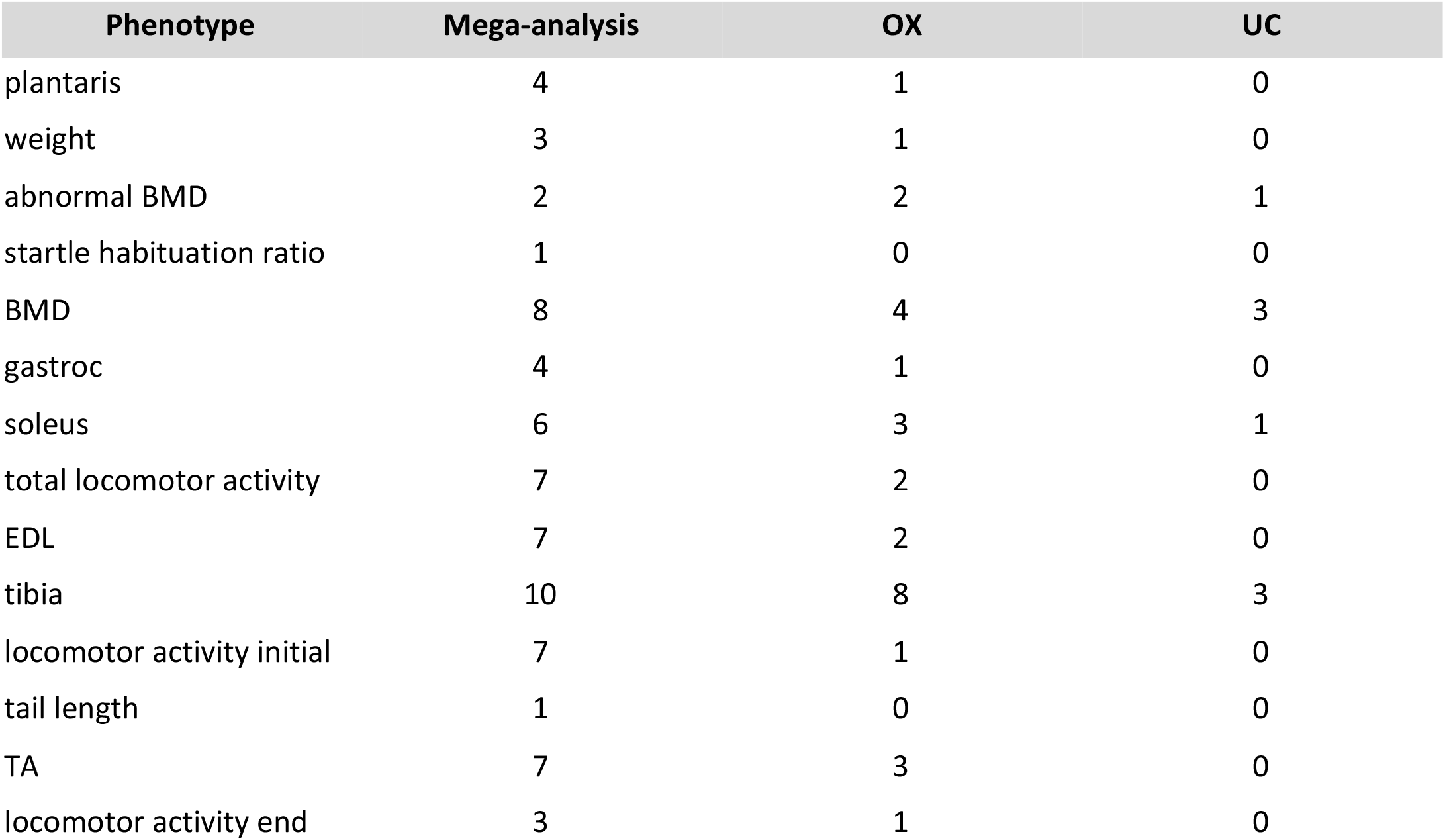
The number of QTLs significant in the mega-analysis (“Mega-analysis”) and the number of the QTLs from the mega-analysis that were also found in each of the component studies (“OX” and “UC”). Phenotype names are defined in Supplemental Table 1

| Phenotype | Mega-analysis | OX | UC |
| --- | --- | --- | --- |
| plantaris | 4 | 1 | 0 |
| weight | 3 | 1 | 0 |
| abnormal BMD | 2 | 2 | 1 |
| startle habituation ratio | 1 | 0 | 0 |
| BMD | 8 | 4 | 3 |
| gastroc | 4 | 1 | 0 |
| soleus | 6 | 3 | 1 |
| total locomotor activity | 7 | 2 | 0 |
| EDL | 7 | 2 | 0 |
| tibia | 10 | 8 | 3 |
| locomotor activity initial | 7 | 1 | 0 |
| tail length | 1 | 0 | 0 |
| TA | 7 | 3 | 0 |
| locomotor activity end | 3 | 1 | 0 |

We explored the contributions of power and confounders (which are presumed to reflect experimental differences between studies) to replication rates in our samples. When power is low, variants that pass the significance threshold are more likely to have inflated effect size estimates, meaning they are less likely to be replicated, a phenomenon that is often called “The Winner’s Curse”. Put another way, when power is low, the positive predictive value falls, so that for a given P-value there will be more false positives.

We applied a statistical framework that jointly models winner’s curse and study-specific heterogeneity due to confounding (Zou et al. 2020). The inputs to this method are the z-scores of the significant variants from the discovery study and the corresponding z-scores from the replication study. The method compares two random effects models: one that accounts for winner’s curse (WC) and another that accounts for both winner’s curse and confounding (WC+C). We compared the expected replication rates under each of these random effects models to the observed replication rates. If the observed replication rate is well explained by the WC model, there is relatively little confounding. However, if the observed replication rate is better explained by the WC+C model, there may be confounding between the studies.

We used the OX cohort as the discovery data set and UC cohort as the replication data set. We jointly modeled Winner’s Curse and study-specific heterogeneity due to confounding in nine phenotypes which had at least three significant loci (model parameters are not robustly estimated with fewer significant loci (Zou et al. 2020)). We computed the expected replication rate under each model for each phenotype and compared these estimates with the observed replication between the two data sets. In order to estimate confidence intervals we applied this method to one hundred randomly divided sets of the OX sample (confounding should not exist when a single study is randomly split in half into two cohorts). To display the results, we plotted the difference between the observed replication and the expected replication after correcting for Winner’s Curse, which should be close to zero when the model explains the observed data.

Figure 5A shows the results modeling Winner’s Curse (WC). The estimate for one phenotype, EDL, overlaps with zero, but in all other cases the Winner’s Curse model predicts higher rates of replication than we observed. The WC+C model jointly models Winner’s Curse and study-specific heterogeneity due to confounding. Taking into account confounds in addition to Winner’s Curse (WC+C) does a better job explaining the observed replication, as shown in Figure 5A. Only one phenotype, bone mineral density, deviates significantly from zero (Figure 5A). In this phenotype, replication was not explained well by either model.

**Figure 5:**
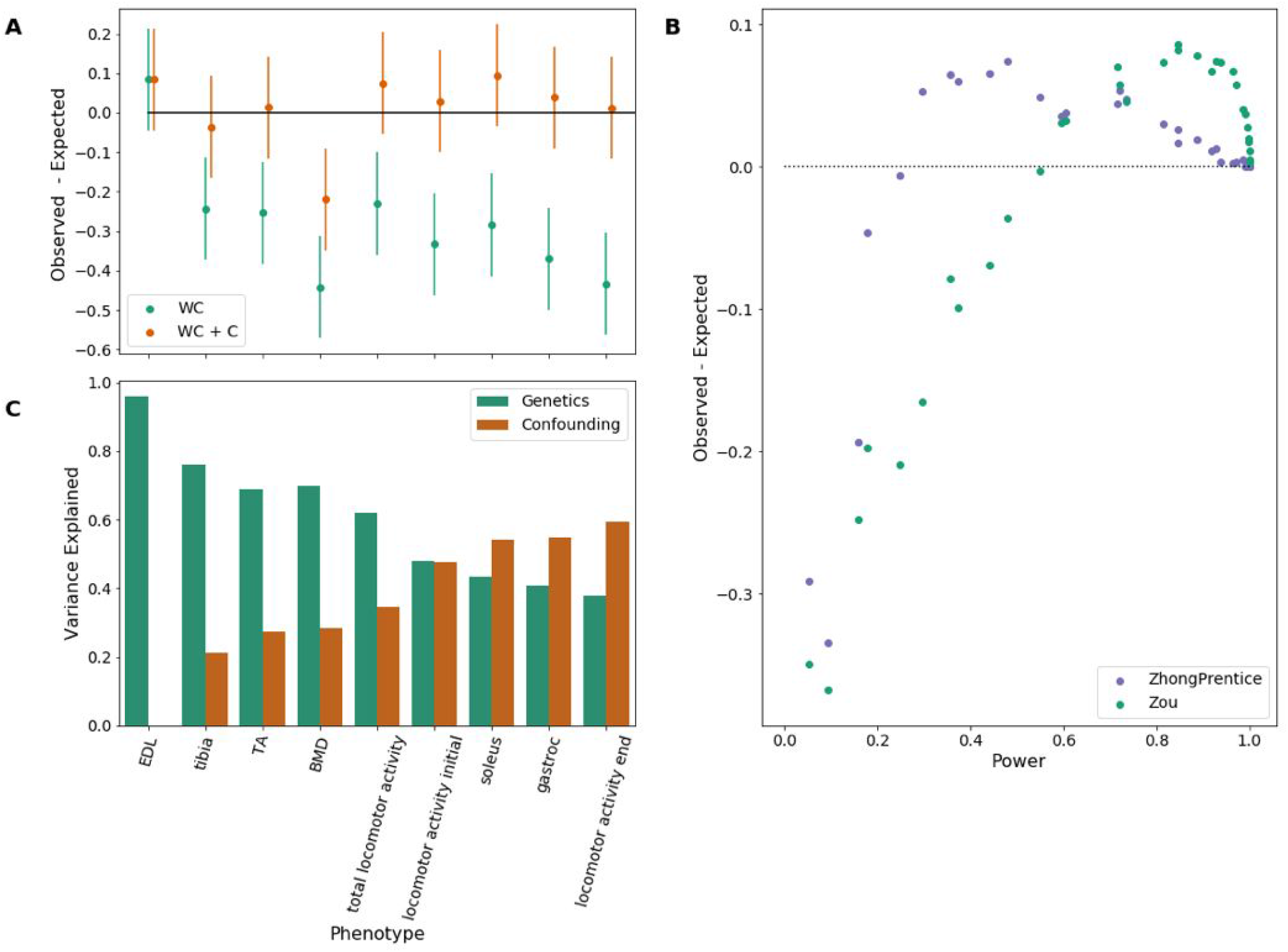
Replication analysis between OX and UC cohorts. A) Difference between the observed and expected replication (y-axis) after accounting for Winner’s Curse (green) and after accounting for both Winner’s Curse and confounding (brown). Empirically computed 95% confidence intervals are shown as vertical bars. B) Simulated summary statistics, with power ranging from 5% to 100% (horizontal axis) are shown analysed for the difference between the observed and expected replication rate (vertical axis). Winner’s Curse was corrected by two methods (blue for Zhong-Prentice (Zhong and Prentice 2008) and green for the Zou WC only method). C) Variance attributable to confounding and Winner’s Curse (genetic effects). The phenotypes are sorted by the amount of variance explained by genetics. There are six phenotypes with at least as much variance explained by genetics as confounding (EDL, tibia, BMD, total locomotor activity, TA, locomotor activity initial) and three phenotypes where the confounding is more substantial (soleus, gastroc, locomotor activity end). Phenotype names are defined in Supplemental Table 1.

We noted that poor replication is observed in more of the behavioral than physiological phenotypes. The former typically have smaller effect sizes relative to physiological traits, as we know from our mega-analysis. Since it has been shown that correction for Winner’s Curse is inaccurate when power is low (Zhong and Prentice 2008; Xiao and Boehnke 2009) we wondered whether the correction we applied was performing appropriately, and whether an alternative correction method might perform better. To address this issue we simulated summary statistics with power levels ranging from 5% to 100% and calculated the difference between estimated and expected replication, using two methods to correct for Winner’s Curse. We used our correction method (Zou et al. 2020) and that of Zhong and Prentice (Zhong and Prentice 2008) and confirmed the presence of bias when power is low (Figure 5B). However, as power increased, the methods performed differently: Zhong-Prentice overestimated replication as power exceeded 20%, while Zou overestimated when power exceeded 50%. The degree of bias in both methods decreased as power approaches 100%. The non-linear relationship of the correction with power, and the different performance of both methods, means that neither provides an optimal solution.

We estimated the relative contribution of Winner’s curse and confounding for each phenotype. Figure 5C shows that the relative contribution varies between phenotypes. Replication rate differences for EDLs depend almost entirely on Winner’s Curse while for soleus, gastroc, and locomotor activity end, confounds outweigh the contribution of Winner’s curse. In general, behavioral phenotypes tended to have more confounding than physiological phenotypes.

We returned to the results of the mega-analysis to categorize the robustness of the findings, based on the results of our replication analysis. Using effect sizes corrected for Winner’s Curse (using (Zhong and Prentice 2008)) we estimated the power in the OX, UC, and mega-analysis studies (Supplemental Table 6). As expected, power was on average higher in the mega-analysis than the component studies (Supplemental Figure 9). We then used these power estimates in combination with replication data and estimates of confounding to divide the mega-analysis results into four categories (Table 3).

**Table 3:**
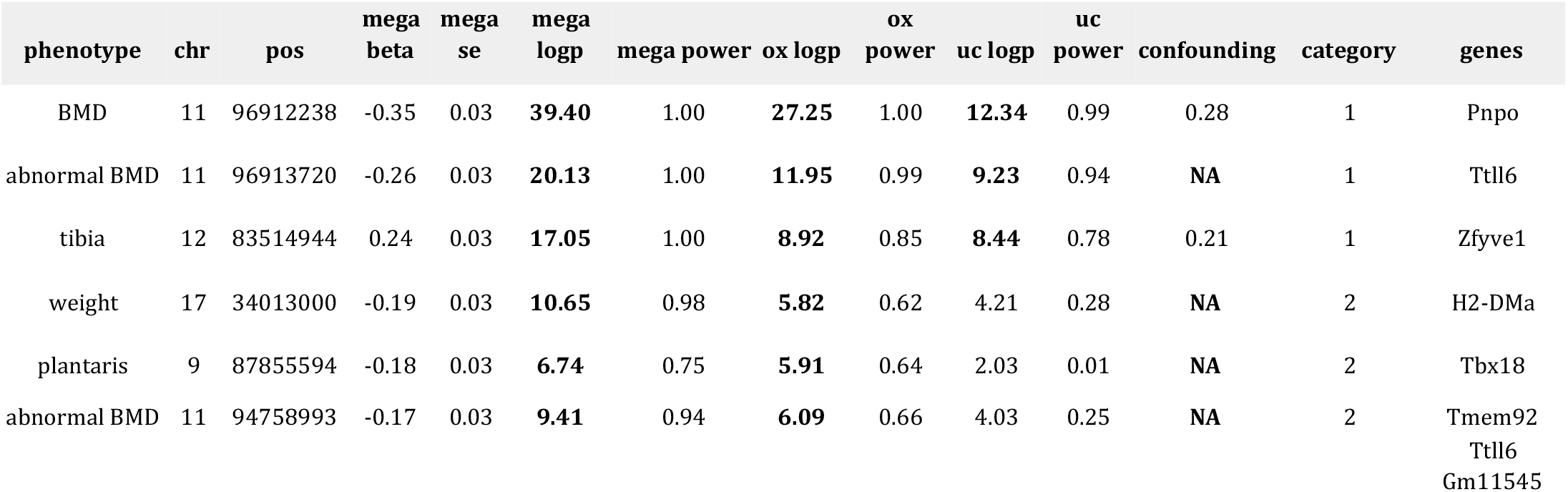
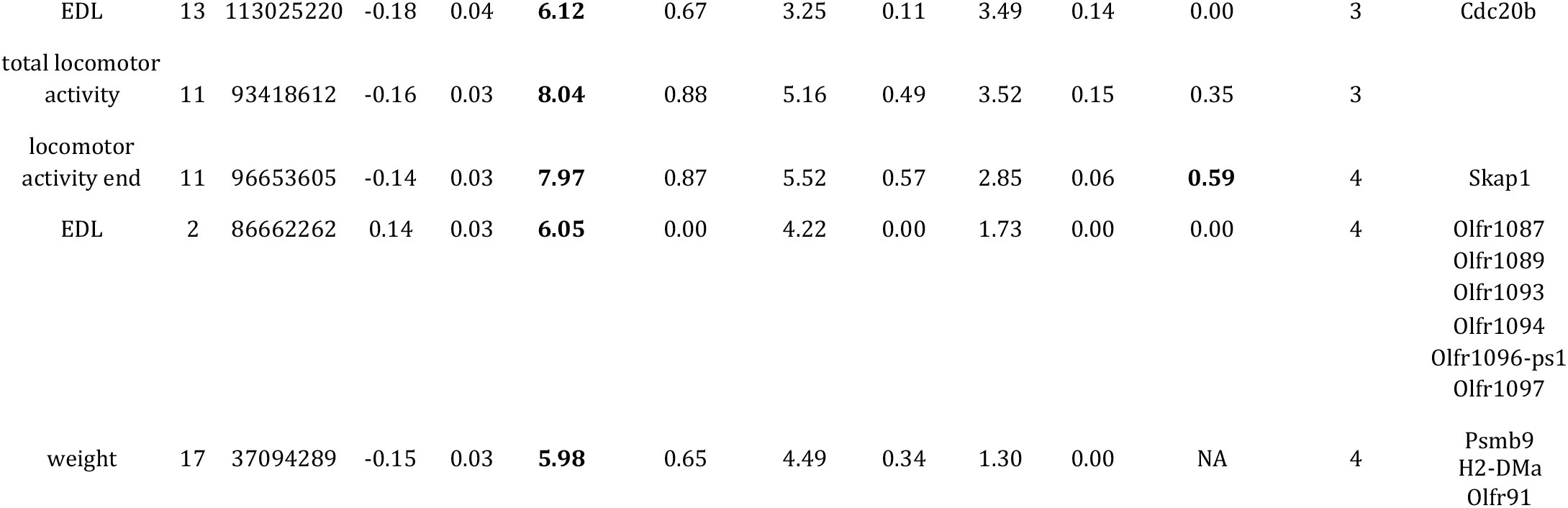
Example QTLs obtained in the mega-analysis, divided into four confidence categories. Phenotype names (“phenotype”) are defined in Supplemental Table 1. The chromosome and position of each QTL are shown in columns “chr” and “pos”. The effect size, standard error, and negative log p-values in the mega-analysis are shown in “mega beta”, “mega se”, and “mega logp”, respectively. Negative log p-values that exceed the genome-wide threshold are bolded. The estimated power in the mega-analysis is shown in “mega power.” The negative log p-values and estimated power for the OX and UC studies are shown in the following 4 columns. The “confounding” column shows the level of confounding observed for each phenotype (not locus-specific), which is the estimated fraction of variance in the GWAS study explained by confounding. Bolded values correspond to phenotypes where the fraction of variance explained by confounding was greater than the fraction of variance explained by genetics (NA indicates that confounding could not be calculated). Categories (“category”) were determined from replication data, estimated level of confounding and estimated power in the mega analysis. Category 1 contains QTLs that were replicated in both the OX and UC cohorts. Category 2 contains QTLs that were replicated in at least one cohort. Category 3 contains variants with low estimated level of confounding. Category 4 contains all other variants. The genes implicated by each of the QTLs are shown (“genes”). See full results in Supplementary Table 6.

The first category consisted of a high confidence set of variants that were significant in the combined study and both component study. 8 of 70 variants in 4 phenotypes were in this first category of variants. The second category consisted of variants that were significant in the combined study and at least one component study. 21 of 70 variants in 12 phenotypes were in this second category of variants. The third category consisted of variants found in phenotypes without evidence of confounding (excluding phenotypes without enough significant variants to determine the confounding level). 14 of 70 variants in 5 phenotypes fell in this category. Finally, the fourth category contained the remaining variants. 27 of 70 variants in 12 phenotypes fell within this category.

## Discussion

By combining phenotypes and genotypes from two independent laboratories, we identified 70 loci for 23 complex traits in a population of 3,076 commercially available outbred mice. The large sample delivered a median QTL interval size of 1.1 Mb. Fine-mapping reduced the number of likely causal variants, with a median reduction in set size of 51%. For all traits analysed our results are consistent with a polygenic architecture, in which the vast majority of causal variants likely lie in non-coding parts of the genome, and with the existence of a considerable degree of pleiotropy, a pattern commonly recognized from GWAS of human complex traits (Visscher et al. 2017). As well as indicating the genetic architecture of the traits, our findings cast some light on the biology of the phenotypes we have mapped, allowed us to examine factors contributing to replication, and hence to assess the robustness of our findings. We discuss these points below.

To assess the robustness of our findings in the mega-analysis, we categorized QTLs by integrating replication data, confounding estimates for each phenotype, and estimates of power in the mega-analysis. We use replication of findings in the OX and UC cohorts to determine the two highest confidence categories. Confidence category 1 consisted of QTLs that replicated in both cohorts. One example QTL in this category was associated with tibia length (chr12:83514944). Through fine-mapping and annotation of variants in the causal set, we implicated the Zfyve1 gene. This gene was also implicated by two other loci associated with gastroc (chr12:83517483) and TA (chr12:83514944) and is expressed highly in these tissues (Lionikas et al. 2012; Lionikas et al. 2013). Zfyve1 is known to bind to Ptdins3P (Lee et al. 2019), which is an important mediator of Akt/PKB kinase activation and upregulation of protein synthesis (Hemmings and Restuccia 2012), providing a potential mechanism for influencing variability in muscle weight. Zfyve1’s link to Ptdins3P along with the GWAS associations with tibia length, gastroc weight and TA weight suggests that Zfyve1 may be involved in modification of the intracellular signal promoting growth.

Confidence category 2 contained QTLs that replicated in one component study but not the other. These associations are less likely to be driven by differences between studies or Winner’s Curse, since they were also significant in a component study with fewer samples than the mega-analysis. For example, chr11:94758993 was associated with abnormal bone mineral density in the mega-analysis and was replicated in the OX study but not the UC study. Through our fine-mapping and nonsynonymous variant annotation, we implicated the gene Tmem92, which has been previously found to be associated with heel bone mineral density in humans in a number of studies (Kemp et al. 2017; Kim 2018; Kichaev et al. 2019; Morris et al. 2019). Additionally, the Tbx18 gene which was implicated by a confidence level 2 QTL for plantaris weight (chr9:87855594), was previously associated with body size phenotypes in human GWAS studies and mouse KO studies (Wu et al. 2013; Lotta et al. 2018; Kichaev et al. 2019; Pulit et al. 2019, Bussen, 2004).

Confidence category 3 contains QTLs that did not replicate in component studies. However, these QTLs were found in phenotypes with relatively lower confounding and had high estimated levels of power (> .5) in the mega-analysis after correcting for Winner’s Curse. A QTL for EDL weight (chr13:113025220) in confidence category 3 implicated gene Cdc20b, which has been previously associated with BMI adjusted waist circumference in human GWAS (Zhu et al. 2020). The estimated power level for this variant was 11% in the OX study, 14% in the UC study and 67% in the mega-analysis. QTLs in confidence level 3 are likely to be true associations that were identified due to the increase in power in the mega-analysis.

In general, we observe that genes implicated using high confidence QTLs are more likely to have support in previously published results. For example, we identified seven olfactory receptor genes from two QTLs associated with EDL weight and body weight that were in the lowest category of confidence (4). Additionally, the EDL QTL (chr2:86662262) had low estimated power after correcting for Winner’s Curse, and the body weight QTL (chr17:37094289) had a high estimated level of confounding between studies. None of the seven genes have been previously associated with related phenotypes in the literature and are unlikely to be true associations. However, there is evidence to support the candidacy of genes at some of the QTLs in this category. For example, a QTL for soleus weight (chr13:9242435) implicates Larp4b, which has been previously associated with atypical femoral fracture and ankle injury in human GWAS (Kim et al. 2017; Kharazmi et al. 2019). Since the soleus is the plantar flexor muscle of the ankle, it is likely to play a role in these two phenotypes. While we estimated substantial amounts of confounding for the locomotor activity end phenotype and the soleus weight phenotype, previous findings may indicate that these genes may have been identified in spite of the confounding we observed.

While these examples highlight genes that have previously been associated with similar phenotypes in mice or humans, other genes (Pnpo, Gm11545, Psmb9, Ttll6) represent novel targets for functional validation in follow-up studies. In particular Pnpo lies at the bone mineral density QTL chr11:96912238, in confidence category 1. The bone mineral density phenotype had relatively low estimated levels of confounding between the OX and UC cohorts, and the power of the mega-analysis after correcting for Winner’s Curse was close to 100%.

Our results also provided us with an opportunity to identify the factors that contribute to replication rates. When we investigated replication between component studies we found that about a fifth (19%) of QTL discovered in OX replicated in UC and just less than a third (32%) of QTL identified in UC replicated in OX. We explored reasons for the low rates of replication and came to the following conclusions.

First, unknown confounds between laboratories limited replication, but this depended on the phenotype. By applying a method that detects the effect of confounding, we observed that for one phenotype there was no impact of confounding (weight of the EDL muscle), while on others the impact was substantial (up to 60% of the variance explained in the case of weight of the plantaris muscle). Some of the differences likely reflect systematic differences between the way the mice were treated at the two testing sites: mice born in Portage, Michigan, were shipped either to Chicago by truck (∼200 Km) or to Oxford by truck and plane (∼6,100 Km). Other confounding factors may include differences in the way the animals were handled or a variety of other laboratory specific differences.

Second, Winner’s Curse contributes to poor replication, even after attempting to correct for its effects. While several strategies have been developed for accounting for Winner’s Curse (Zhong and Prentice 2008; Sun et al. 2011a; Xu et al. 2011; Ferguson et al. 2013; Huang et al. 2018) we lack rigorous comparisons of their relative performance. Here, we found that when power is less than 20%, correction methods underestimate the impact, and at power greater than 50% over-correction occurs. In other words, the relationship between power and bias due to Winner’s Curse depended on the method used. The importance of this observation is that since power is rarely accurately estimated, it is difficult to accurately assess the bias attributable to Winner’s Curse.

Current human GWAS usually report results only from their meta-analyses, following the argument that joint analysis is more powerful than replication-based analyses (Skol et al. 2006), and rarely rely on replication as a touchstone for the dependability of their findings. Results obtained in this way are usually regarded as robust (Marigorta et al. 2018). However, replication rates may be inconsistent with the P-values and effect sizes reported in discovery cohorts; for example a replication rate of 48% was found in a survey of 100 papers (Palmer and Pe’er 2017). ‘Winner’s Curse’ was invoked to explain the disparity (Palmer and Pe’er 2017), but, as we have seen, the performance of corrections for the bias depends on the power of the study and the correction method used, making it impossible to assess the true contribution of Winner’s Curse.

The bulk of human GWAS findings since 2008 are replications of associations that have been described at least twice (Marigorta et al. 2018), thus allaying concerns that the findings are not robust. The same cannot be said for mouse GWAS, where it is rare for a trait to be mapped multiple times at all, much less to be mapped multiple times in the same population. Furthermore, the genetic structure of mouse mapping populations is quite unlike that of outbred human populations. For example, no human population consists of fully inbred individuals, a commonly used design in mouse experiments (as in the recombinant inbred strains of the collaborative cross, or the inbred strains that constitute the hybrid mouse diversity panel (Flint and Eskin 2012)). Replication across such different populations is more difficult than within a population (as already documented for human studies (Palmer and Pe’er 2017)). We show here that obtaining robust results in mice is demanding because of difficulties in obtaining a replication population, the impact of Winner’s curse and phenotype specific confounds. As a first step we derive categories of confidence to our mapping results by combining information about replication, confounding, and power. We thus categorized loci into one of four confidence levels. Our approach helps alleviate concerns about reproducibility (Button et al. 2013), but needs further development to produce a systematic method to evaluate confidence quantitatively, rather than categorically as we have attempted here. This framework may also be applicable to human GWAS especially when replicating in admixed populations, when phenotyping may be difficult.

## Methods

### Subjects

All animals originated from the same colony of outbred mice, the Crl:CFW(SW)-US_P08 stock, maintained by Charles River Laboratories in Portage, Michigan at the time of the study. Details about the OX and UC mice are described elsewhere (Nicod et al. 2016; Parker et al. 2016). The OX mice were purchased between December 2009 and September 2011 at 4-7 weeks of age and phenotyping started at 16 weeks. The UC mice arrived at 7 weeks of age between August 2011 and December 2012, and testing started after a 2 week acclimatisation period. Both the OX and UC mice were maintained on a standard 12:12h light-dark cycle with water and standard lab chow available *ad libitum* and housed 3 (OX) or 4 (UC) per cage.

### Phenotypes

The effect of covariates, including sex, weight and batch, was tested in each dataset separately (OX and UC) and introduced in a linear model to calculate residuals when the effect was significant (weight was retained for mapping body weight). OX and UC residuals were then quantile-normalised before being merged in a single dataset used for the genetic analysis. Supplemental Data 1 lists the mean and standard deviation of all measures obtained in both studies and the covariates included in the linear model to calculate the residuals. Many of the behavioral phenotypes were not independent. For example, there were multiple phenotypes within the category of locomotor activity. They represent measures related to locomotor activity at different timepoints during the testing period (eg. 0-15 min, 15-30 min, etc.) as well as summary data representing changes in activity over time (eg. activity decay) that are likely correlated. Descriptions of all phenotypes can be found in Supplemental Table 1. The following section describes methodological differences between OX and UC. Further details of methods applied in the individual studies can be found in separate publications.

#### Locomotor Activity

In both centres the baseline activity of the mice was recorded during the first 30 minutes of testing. The activity of the OX mice was recorded at 16 weeks after placement into a new plastic home cage (46 cm x 15 cm x 21 cm) using a photoactivity system from San Diego Instruments (San Diego, CA) which has seven infrared photobeams crossing the width of the cage floor.

The UC mice, ∼51 days of age at the time of test (SD = 4), were injected i.p. with physiological saline (0.01 ml/g body weight) immediately prior to the test and placed in the center of a OF chamber (AccuScan Instruments, Columbus, OH) consisting of a clear acrylic arena (40 cm x 40 cm x 30 cm) placed inside a frame containing evenly spaced infrared photobeams.

Four phenotypes were analysed for this study: locomotor activity initial (0-15 min), locomotor activity end (15-30 min), locomotor activity total (0-15 min), and activity decay. Activity decay was defined as the decrease in activity from beginning to end.

#### Conditioned Fear

In OX, conditioned fear was tested over two days in four San Diego Instruments chambers with mice at 17 weeks of age. On the first day of the test mice were subjected to a 13 minute training session during which they received two electric foot shocks (0.3mA, 0.5sec) preceded by a 30 second tone. In the morning of the second day of the test the mice were placed in the same enclosure for five minutes and fear associated with the context was measured by the amount of freezing. In the afternoon the animals were placed in a different enclosure for five minutes where they were subjected to two 30 second tones without any paired electric shock. Freezing behavior during all sessions of the test was scored using a VideoTrack automated system (Viewpoint, Champagne Au Mont D’Or, France).

UC mice were tested for CF at ∼ 63 days of age (SD = 2.9) over three consecutive days, each consisting of a seven minute trial: on the first test day, mice were conditioned to associate a test chamber and a tone (85dB, 3kHz tone lasting 30 seconds) with a shock (2-second, 0.5-mA foot shock delivered four times); on the second test day, the mice were re-exposed to the same context, but no tones or shocks were given; on the third day, mice were exposed to the conditioned stimulus (the tones), but in a different environment. Mice were tested in four chambers obtained from Med Associates (St. Albans, VT, USA) and immobility, or “freezing” behavior, was recorded by analyzing digital video with Freeze Frame software (Actimetrics, Evanston, IL, USA).

Phenotypes from the conditioned fear tests consisted of eight measurements of immobility collected in OX and UC mice. On day one we measured average proportion of freezing during the pre-training interval (UC: 30-180 seconds, OX: 180-355 seconds) before exposure to tones and shocks (baseline freezing D1), as well as the average proportion of freezing during exposure to the first conditioned stimulus corrected for baseline freezing (corrected freezing to tone alone). On day two we measured freezing during the context test corrected for baseline (corrected freezing to context). On day three, we measured the average proportion of time freezing in the altered setting during the 30-second intervals in which the tones were presented corrected for baseline (corrected freezing to cue).

#### Prepulse Inhibition

OX mice were tested at ∼17 weeks of age following a different protocol (Yee et al. 2005)combining exposure to loud pulses of three different intensities (100, 110 and 120 dB), also preceded by 3-12dB prepulses. For this analysis, only measures of startle elicited by the 120dB pulse (similar to UC) were considered.

UC mice were tested at a mean age of 74.5 days (SD = 2.2). During PPI, the mice were exposed to loud pulses (120 dB) that caused them to exhibit the startle response. Occasionally, the exposure to the loud pulse was preceded by a barely perceptible “prepulse” (3–12 dB over background levels), which inhibited the startle response to varying extents. The UC PPI testing procedures follow protocols detailed in previous papers (Palmer et al. 2000; Palmer and Airey 2003; Palmer et al. 2004; Shanahan et al. 2009).

Both OX and UC Prepulse inhibition tests were performed in chambers and apparatus from the same manufacturer and model (San Diego Instruments, San Diego, CA, USA), capturing mouse movement using a piezoelectric accelerometer, then converted into digital data and recorded on a computer. Before the start of each test day, the apparatuses were calibrated according to the manufacturer’s instructions.

Five phenotypes were analysed for this study: startle habituation difference, startle habituation ratio, startle, PPI with +6db prepulse and PPI with + 12db prepulse. In UC, startle habituation difference was defined as the average startle amplitude during the fourth pulse-alone trials subtracted from the average startle amplitude during the first pulse-alone trials. In OX this was calculated by subtracting the last block from the first block for the 120 dB stimulus. In UC, startle habituation ratio was defined as the average startle amplitude during the first pulse-alone trials divided by the average startle amplitude during the fourth pulse-alone trials. In OX this was calculated by dividing the first block for the 120 dB stimulus by the last block. Startle was defined as the average startle amplitude to the 120 db pulses during the second and third blocks. PPI with +6db prepulse was defined as the average pre-pulse inhibition during blocks 1 and 2 to the 6 dB prepulse. PPI with +12db prepulse was defined as the average pre-pulse inhibition during blocks 1 and 2 to the 12 dB prepulse.

#### Musculoskeletal Traits

Weight of hind limb muscles and length of tibia were collected by the same experimenter for both the OX and UC mice. Following sacrifice, at ∼20 weeks for the OX mice and ∼90 days of age for the UC mice (M = 91.2, SD = 2.6), one leg was cut off just below pelvis, tubed and transferred into a −70 C freezer, then shipped on dry ice to Dr. Arimantas Lionikas at the University of Aberdeen. On the day of dissection, the leg was defrosted and two dorsiflexors, tibialis anterior (TA) and extensor digitorum longus (EDL), and three plantar flexors, gastrocnemius (gastroc), plantaris and soleus, were removed under a dissection microscope and weighed to a precision of 0.1 mg on a balance (Pioneer, Ohaus). Then, the soft tissues were stripped off from the tibia, and the length of the tibia was measured to a precision of 0.01 mm with a digital caliper (Z22855, OWIM GmbH & Co).

#### Bone Mineral

In the OX mice, bone mineral content of the tibia was measured with the Faxitron MX-20 scanner (Faxitron Bioptics LLC, AZ, USA) using methods adapted from (Bassett et al. 2012). ImageJ (V1.48p, National Institutes of Health, USA) was used to quantify the apparent bone mineral content and the mean of the value obtained for the entire bone used for analysis. In the UC mice, bone mineral density for the entire isolated femur was assessed by Dual X-ray absorptiometry (DXA) using a GE-Lunar PIXImus II Densitometer (GE-Lunar, Madison, WI). To admit a normal distribution, we transformed the BM measurements, which were ratios, to the (base 10) log-scale. Lastly, we measured abnormal BMD, which is a dichotomized version of the BMD phenotype, and the only phenotype collected that was not quantitative.

#### Body Weight and Tail Length

Body weight was measured at ∼17 weeks in OX mice and at ∼13 weeks of age (91.2, SD = 2.6) in UC mice. Tail length (in cm) was measured at the time of sacrifice, as the distance from the base of the tail to the tip of the tail (OX: ∼20 weeks of age, UC: same as body weight measurement).

### Genotypes

Sample BAM pre-processing including mapping and quality control is as previously described (Nicod et al. 2016; Parker et al. 2016). Reads were aligned to the reference genome assembly (NCBI release 38, mm10). Imputation was carried out using STITCH (Davies et al. 2016) (version = 1.6.0, K = 4, nGen = 100) at 7.07 M bi-allelic autosomal and chromosome X SNPs on the combined set of N = 2073 Oxford + N = 1161 Chicago mice. No samples were removed post-imputation. Genotypes for assessing imputation accuracy were generated on the MegaMUGA array (Collaborative cross consortium 2012) for 48 Oxford and 48 Chicago samples at 77,588 SNPs. After applying standard per-sample (minimum quality scores, removal of samples failing sex checks) and per-SNP (missing less than 5%, Hardy-Weinberg Equilibrium (HWE), p-value greater than 1×10^−10^) quality thresholds, genotypes for 44 Oxford and 32 Chicago samples at 73,218 SNPs were available for accuracy assessment. Accuracy was assessed using correlation (r^2^) SNP-wise between imputed and true (array) genotypes. Post-imputation SNP QC was carried out for each SNP using each of the combined set of all samples, the Oxford samples only, and the Chicago samples only. Metrics of INFO score and HWE p-value (using hard genotype calls where the maximum posterior genotype probability was greater than 0.9) were calculated for each. Array genotypes were intersected with the imputed genotypes (14,942 SNPs), and only sites with a minimum allele frequency of 1% across each of the three sets kept (13,933 SNPs). The 3.15 M SNPs from STITCH were pruned for linkage disequilibrium using PLINK (Purcell et al. 2007). The genotype dosages were read into PLINK, and using PLINK’s hard calling threshold, were converted to genotypes. Using these genotypes, the SNPs were filtered for the LD measure, r^2^. Jointly calling the OX and UC samples in this manner generate a high confidence set of SNPs that was different from those used in their prior studies (Nicod et al and Parker et al).

### Mapping of traits in the combined set

The genotype likelihoods at 3.15 M SNPs, obtained using STITCH, were used to perform associated studies in the joint set of Oxford and Chicago samples. The genotype likelihoods were converted to dosages of the alternate allele. Using the dosages for the combined set of samples, association analyses were performed using linear mixed models in GEMMA (v0.96) (Zhou and Stephens 2012) with a pre-calculated genetic relationship matrix (GRM) also from GEMMA. Phenotypes were transformed tby quantile normalization after removing the effects of age and sex, all analyses performed using the R progamming language (R Development Core Team 2010). We did not quantile normalize within sex because sex was used as a covariate to generate the residuals, thus removing the difference in the means. In addition to performing the association analyses on the combined set of Oxford and Chicago samples, association analyses was also performed on the individual sample sets (OX and UC). Finally, the Oxford sample set was randomly split into two equal sized samples sets 100 times. Association analyses were performed on these Oxford subsets as well. Independent QTLs were obtained using a peak caller and a threshold of 0.6, where the QTLs with the highest log p-values were retained.

### Computation of significance thresholds for association analyses

The significance thresholds for the p-values from the linear mixed model association analyses were computed in two different permutation approaches. First, we used a “naïve” permutation approach (Cheng and Palmer 2013). In this approach, for each phenotype, we permuted the phenotype values across the combined set of samples. Then the association analysis was repeated on the permuted phenotypes, and the most significant p-value was retained. For each phenotype, the naïve permutation analysis was repeated 100 times, resulting in a total of 3400 most significant p-values. The permutation threshold was computed as the 5^th^ percentile of the distribution of the most significant p-values from across the all phenotypes; combining the phenotypes was justified because all phenotypes had been quantile normalized, therefore they had identical distributions.. The same permutation threshold was then used for all phenotypes. Separate p-value permutation thresholds were computed for the different sample sets: combined, OX, UC and the two Oxford subsets (OX1 and OX2).

We compared the naïve permutation method to using another permutation approach - henceforth referred to as the “decorrelated” method (Abney 2015). First, the centered GRM was computed using the genotype dosages from all the chromosomes. Then the inverse square root of the GRM was computed. The phenotype and genotype vectors are pre-multiplied by the inverse square root of the GRM, resulting in decorrelated measurements. Since the phenotypes and the genotypes are decorrelated, a linear model is used instead of a linear mixed model, resulting in faster computation times. For each phenotype, 100 permutation replicates were performed by permuting the decorrelated phenotypes and performing association analyses using linear models. Similar to the naïve approach, the most-significant p-value was retained from each permutation replicate across the 3400 phenotype-replicate combinations. Again, the significance threshold was computed as the 5th percentile of the distribution of the most-significant p-values. The significance thresholds obtained using the decorrelated method were close to the naive permutation method (Supplemental Figure 4). Since the decorrelated method was less computationally intensive, and the two approaches led to very similar significance thresholds, we use the thresholds obtained using the decorrelated method for our results.

### Genetic correlation

We constructed local linkage disequilibrium (LD) weighted genetic relatedness matrices (GRMs) using imputed dosages at the common set of SNPs for Oxford and Chicago studies separately, and a combined GRM for all mice in both studies, using LDAK version 5.9 (Speed et al. 2012). We then inferred “narrow-sense” heritability from genome-wide SNPs (h2) at 23 phenotypes measured in both studies, and genetic correlations (rG) between the two component studies in phenotypes with non-zero heritability. Estimates for h2 at each phenotype were obtained using the individual study GRMs using restricted maximum likelihood (REML) implemented in LDAK, while rG for each phenotype was estimated with the combined GRM using bivariate REML implemented in GCTA version 1.93.2 (Yang et al. 2011). All h2 and rG calculations were made with PCs 1 to 20 from PCA on the respective GRMs included as fixed effect covariates. We performed a mega-analysis for all phenotypes with significantly non-zero rG (p<0.05).

### Confidence intervals and fine-mapping

Confidence intervals were estimated by simulation. At each QTL, a residual phenotype was constructed by removing the effect of the top SNP at the QTL from the phenotype vector used in the QTL mapping above. This abolished the effect attributable to the QTL whilst maintaining genetic contributions from elsewhere in the genome. One thousand SNPs were then randomly selected, subject to the constraint that they were within 2.5Mb of the top SNP and were polymorphic in the subset of individuals phenotyped for the trait. A causal variant was simulated at the SNP, with an effect size matching that of the top SNP, taking account of the allele frequency, and added to the residual phenotype. A local scan of the region using the same mixed model but the simulated phenotype was performed and the location and logP of the simulated top SNP recorded. Across the 1,000 simulations, we estimated the distribution of the drop ∆ in logP between the simulated top SNP and the simulated causal SNP (this was zero when the top and causal SNPs coincided). We used the fraction of simulations f(∆) within ∆ to determine confidence intervals for the original phenotype data. Thus we identified the range of SNPs within 1.5Mb of the top SNP and with a logP drop less than ∆ to define the 100f(∆)% confidence interval for the QTL. We used the pruned SNPs within the 95% confidence intervals as input to the SusieR fine-mapping method. We ran the SusieR method (Wang et al. 2020) using the susie_rss function with L=1 and coverage=.95. We annotated all variants within the 95% causal sets and all variants in perfect LD using ANNOVAR (Wang et al. 2010).

### Modeling Winner’s Curse and confounding

We applied two random effects models proposed in (Zou et al. 2020). The z-scores of the significant variants in the discovery study and the corresponding z-scores in the replication study are used as input to learn the three unknown parameters (σ_*g*_^2^, σ_*c*1_^2^, σ_*c*2_^2^) using maximum likelihood. σ_*g*_^2^ is the variance in the true effect size. σ_*c*1_^2^ and σ_*c*2_^2^ are the variances in the study-specific confounding effects for the discovery and replication studies respectively. The first model (WC) only corrects for Winner’s Curse and assumes that the discovery and replication studies have a shared genetic effect (λ ∼ *N* (0, σ_*g*_ ^2^)). This model corrects for Winner’s Curse by modeling the conditional distribution of the replication statistics given discovery statistics 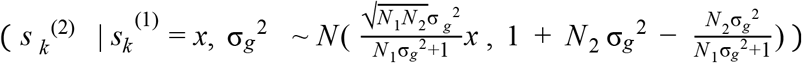, where *N* _1_ and *N* _2_ are the sample sizes of the discovery and replication studies respectively and *k* is a genetic locus used as input to the method). The second (WC+C) corrects for both Winner’s Curse and study-specific confounding effects for the discovery (δ^(1)^) and replication studies (δ^(2)^), where δ^(1)^ ∼ *N* (0, σ_*c*1_^2^) and δ^(2)^ ∼ *N* (0, σ_*c*2_^2^). This model corrects for both Winner’s Curse and confounding using a similar conditional distribution 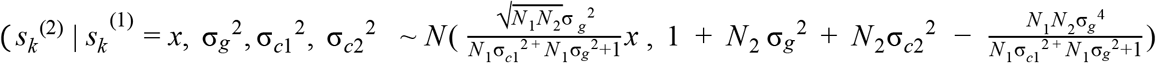, where x is the value of the z-score in the discovery study). We apply this method only to the 9 phenotypes with at least 3 significant loci.

We compared the expected replication rate under the two models to the observed replication. The observed replication was computed as the fraction of variants that were significant in discovery study that were also nominally significant and having the same sign in the replication study. The expected replication rate (*r*) under each model was computed as 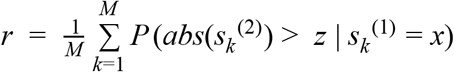, where M is the number of variants significant in the discovery study, and z is the significance threshold for the replication study.

### Winner’s curse simulations

To compare the accuracy of the Winner’s Curse corrections., we simulated 1000 z-scores for the discovery and replication studies using a range of effect sizes (λ ∈ [.1, .11, .12, .13, .14, .15, .16, .17, .18, .19, .2]) and sample sizes (*n* ∈ [1000, 1500, 2000]) by drawing independently from normal distributions 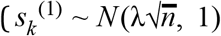 and 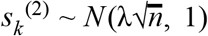 for SNP *k*). We simulated every combination of effect size and sample size in order to simulate a range of power levels. We then used a Bonferroni corrected significance threshold of 5e^-4^ to identify variants significant in discovery study. We considered variants with the same sign in the replication study meeting a nominal significance threshold to be replicated. We computed the true replication rate as the number of replicated variants divided by the total number of significant variants in the discovery study. We then used the simulated data as input to the WC method in (Zou et al. 2020) and (Zhong and Prentice 2008)to correct for Winner’s Curse. We estimated the estimated replication rate under the WC model in (Zou et al. 2020) and compared this value to the true replication rate. After correcting the summary statistics using (Zhong and Prentice 2008), we computed the replication rate after correcting for Winner’s Curse and compared this with the true replication rate.

### Power calculations

We estimated the power of the association statistics obtained from the OX, UC, and combined data sets using (Zhong and Prentice 2008) to correct for Winner’s Curse. This method corrects for the bias from Winner’s Curse and provides an estimate of the true effect size (β_*true*_) as follows.

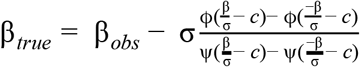

Where β_*obs*_ is the observed effect size, o is the standard error, *c* is the significance threshold, ϕ (*x*) is the standard normal density function, and ψ(*x*) is the standard normal cumulative density function. We use the estimates of the true effect size to compute the power as the probability of observing a significant result under a normal distribution 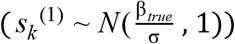.

## DATA ACCESS

The [specify data types] data generated in this study have been submitted to the NCBI BioProject database (https://www.ncbi.nlm.nih.gov/bioproject/) under accession number XXX (including public database accession numbers for all newly generated data and/or reviewer links to deposited data when accessions are not yet public. Previously published accessions should be included in the Methods section where appropriate),

## ACKNOWLEDGMENTS

This work was partially supported by National Institutes of Health grants (R01MH115979 (JF), and R01GM097737 (AAP) and P50DA037844 (AAP)). J.Z is supported by a National Science Foundation Graduate Research Fellowship under Grant DGE-1650604. Publication charges for this article have been funded by 1R01MH115979.

## Author contributions

J.F., A.A.P., and R.M conceived the study. J.Z., J.F. and S.G performed the bioinformatics analysis. C.P. and J.N. prepared the phenotypes. R.W.D generated the genotypes. J.Z., C.P., S.G, N.C, A.L. A.A.P and J.F. wrote the manuscript. All authors read and approved the final manuscript.

