## Supplemental Materials for "Analysis of independent cohorts of outbred CFW mice reveals novel loci for behavioral and physiological traits and identifies factors determining reproducibility"

### Table of Contents

Supplemental Figures 2

Supplemental Tables 12

Supplemental Data 16

### Supplemental Figures

*Supplemental Figure 1: Pairwise phenotypic correlations. Phenotypes have the same order on the horizontal and vertical axes. The color of the phenotype label corresponds to whether it is a physiological (blue) or behavioral (red) phenotype. The color corresponds to the Pearson correlation between each pair of phenotypes. Phenotypes within the same category are more correlated to each other than phenotypes in different categories.*

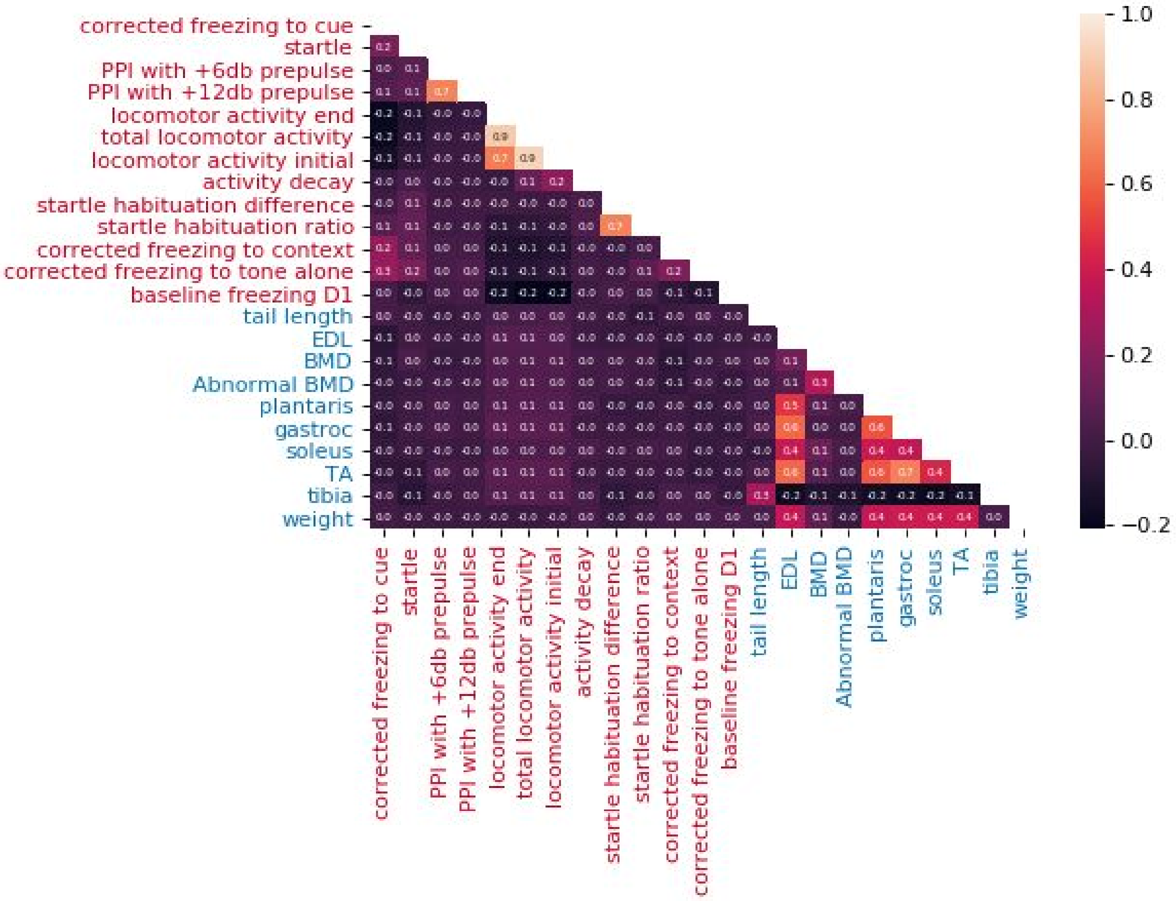

*Supplemental Figures 2A and 2B*

*Shown are the discriminatory capabilities of INFO (shown in (A)) and HWE (shown in (B)) scores. Columns are results using scores calculated from either only Oxford samples, only Chicago samples, or all samples. Rows are as follows. First row, histogram. Second row, one minus empirical cumulative density function. Third row, scatter plot of per-SNP accuracy vs quality control measure, with Chicago SNPs plotted second. Fourth row, windowed accuracy per bin for measure i.e. only SNPs with that value of the measure. Fifth row, accuracy for all SNPs whose value on the measure was greater than or equal to the measure value. (C) Quantile-quantile plot for total locomotor activity phenotype. The theoretical quantiles of the negative logarithm p-value distribution (horizontal axis) are plotted against the observed distribution (vertical axis). The colors correspond to the mega-analysis (blue), OX (orange), and UC (green) cohorts.*

A.

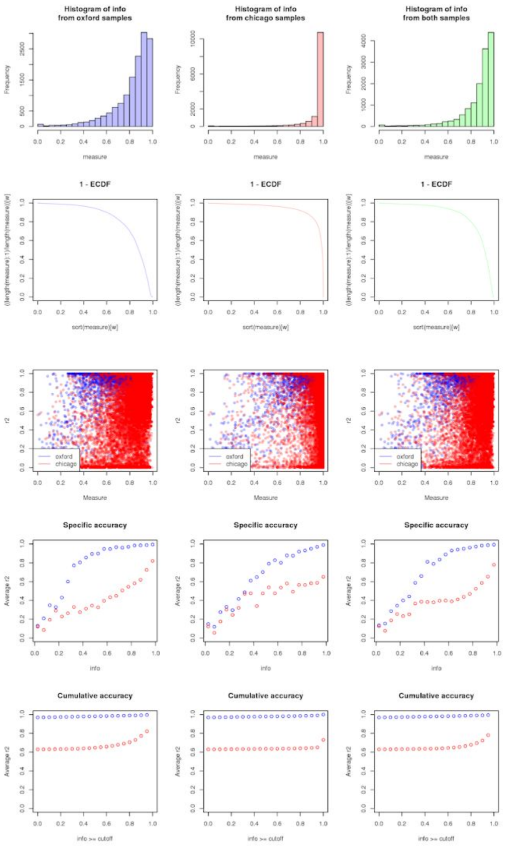
Histogram of info
from oxford samples
Histogram of info
from chicago samples
Histogram of info
from both samples
Frequency
measure
1 - ECDF
sort(measure)[w]
Measure
r2
oxford
chicago
Specific accuracy
Average r2
info
Cumulative accuracy
info >= cutoff

B.

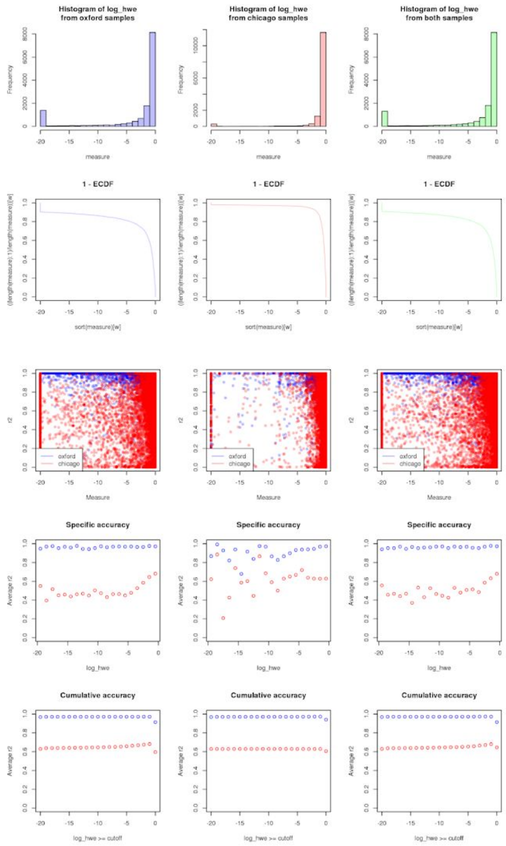
Histogram of log_hwe
from oxford samples
Histogram of log_hwe
from chicago samples
Histogram of log_hwe
from both samples
Frequency
measure
1 - ECDF
sort(measure)[w]
Measure
r2
oxford
chicago
Specific accuracy
Average r2
log_hwe
Cumulative accuracy
log_hwe >= cutoff

C.

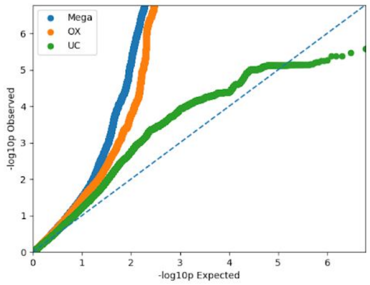
Mega
OX
UC
-log10p Observed
-log10p Expected

*Supplemental Figure 3: Genetic correlation between all phenotypes of OX1 and OX2 data sets.. Each dot represents the estimated genetic correlation, and the error bars show the 95% confidence intervals obtained using the standard errors.*

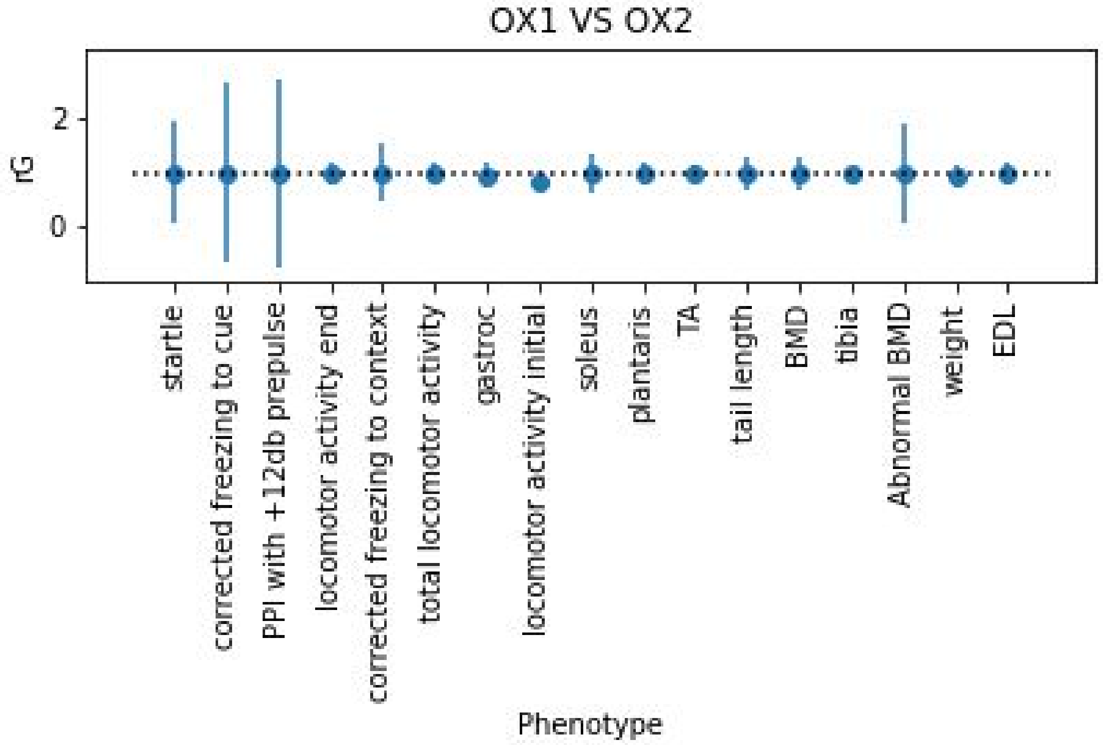

*Supplemental Figure 4: Comparison of significance thresholds using decorrelated and naïve methods. We applied permutation-based methods ("decorrelated" and "naïve") to identify significance thresholds in the three cohorts (OX - red, UC - blue, Combined- green). The thresholds obtained using the decorrelated method (solid lines) tend to be more stringent than the thresholds obtained using the naïve method (dotted lines). We use the decorrelated thresholds in our analysis (* $\alpha = .05$ *).*

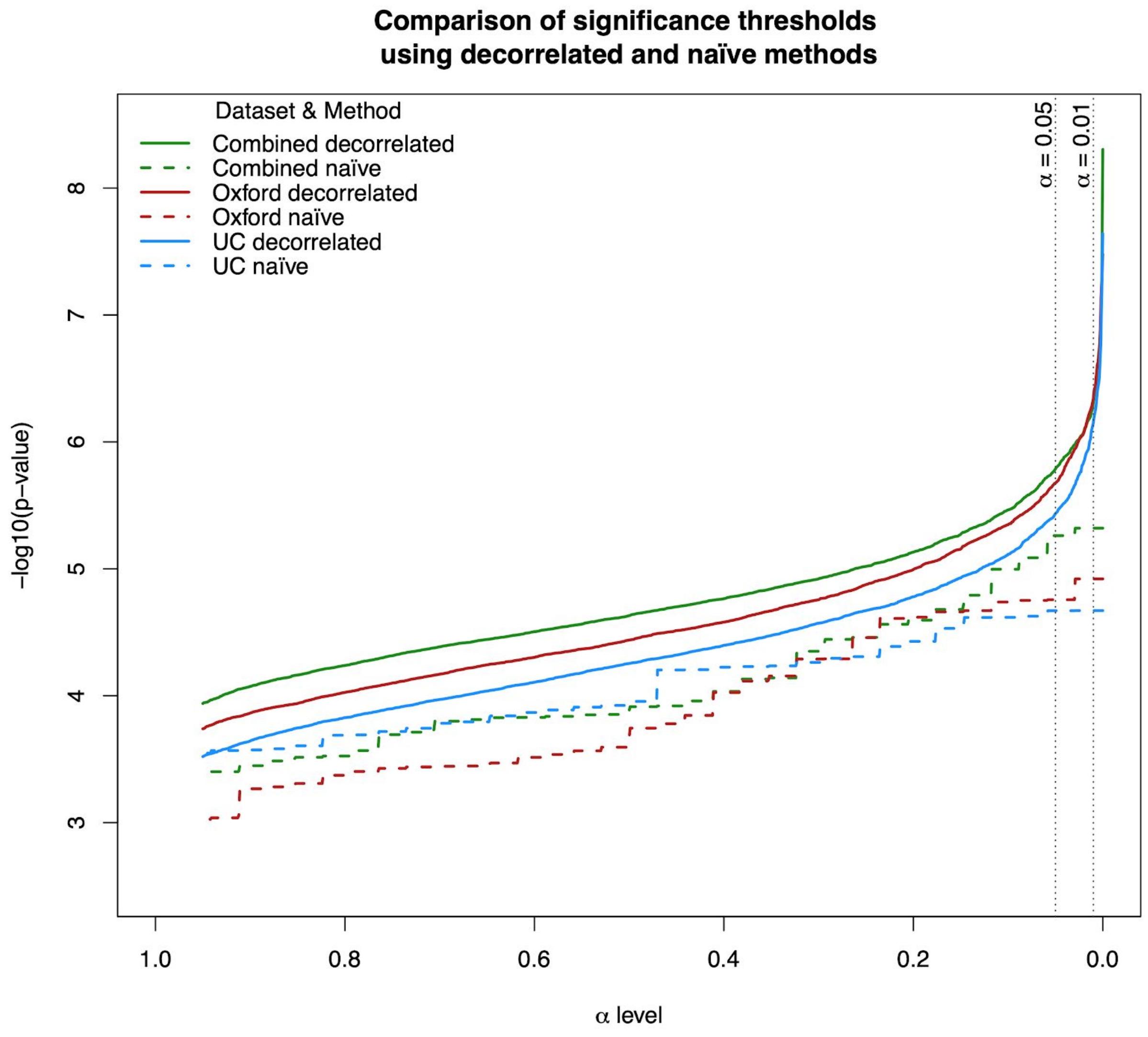

*Supplemental Figure 5: Distribution of the widths of the confidence intervals for the mega-analysis QTLs. The median confidence interval size is 1.1 MB.*

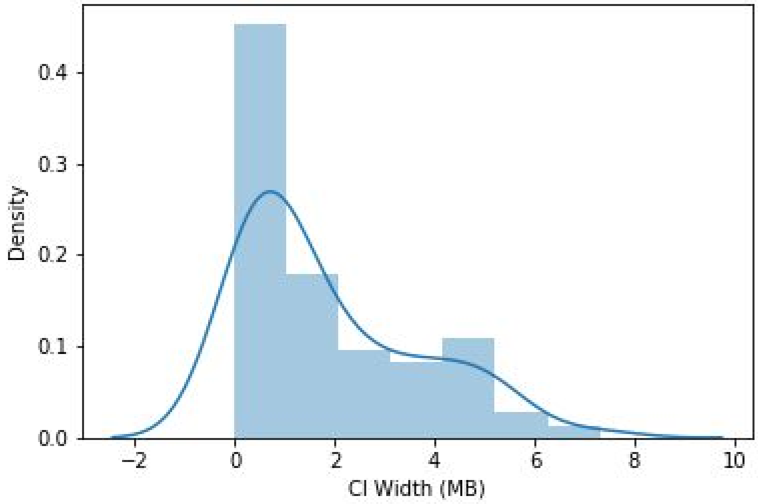

*Supplemental Figure 6: We used all variants within the 95% confidence intervals of the lead SNPs as input to the SusieR fine-mapping framework. The number of SNPs used as input to the fine mapping ("Finemapping Input") is on average larger than the number of SNPs output in the credible sets ("Credible Sets"). The median reduction in the number of SNPs from the fine mapping input to the credible sets was 51%.*

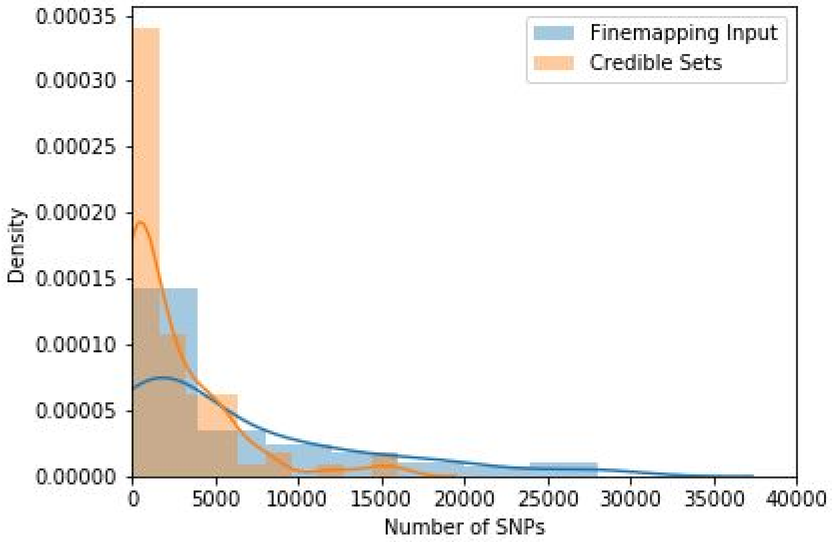

Supplemental Figure 7 : *Co-localization of QTLs for different traits. The horizontal axis lists the QTLs, the vertical axis gives the negative logarithm (base 10) of the association P-value. The color of the text corresponds to the phenotype group of the QTL, with red corresponding to behavior and blue to physiology. The vertical axis shows the negative logarithm p-values of association statistics, where black diamonds correspond to the value of the QTLs. Circles represent the values at other traits, where red corresponds to behavior and blue to physiology. In some cases, behavior QTLs colocalize with physiology QTLs and vice versa. The horizontal black line corresponds to a 5 percent Bonferroni threshold (p< 2.1e-05) that corrects for the number of QTLs and phenotypes tested.*

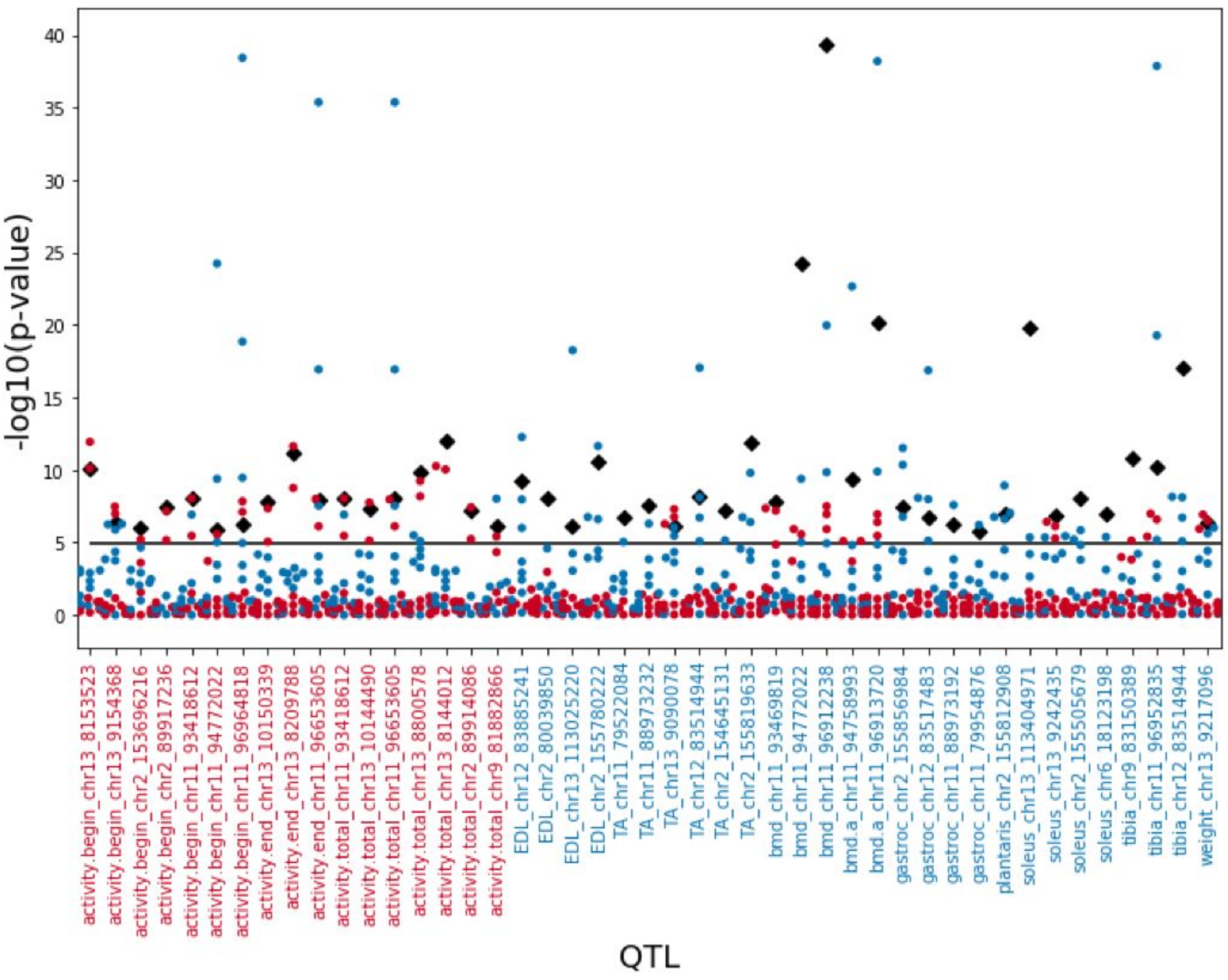

*Supplemental Figure 8: Genome-wide representation of QTLs identified for all phenotypes in the two studies, OX and UC. Light grey dots show association for the measures where a QTL was detected. The most significant SNPs at each QTL are marked with a colored dot, where blue represents physiological and red behavioral traits. Some peaks contain multiple QTLs from related traits.*

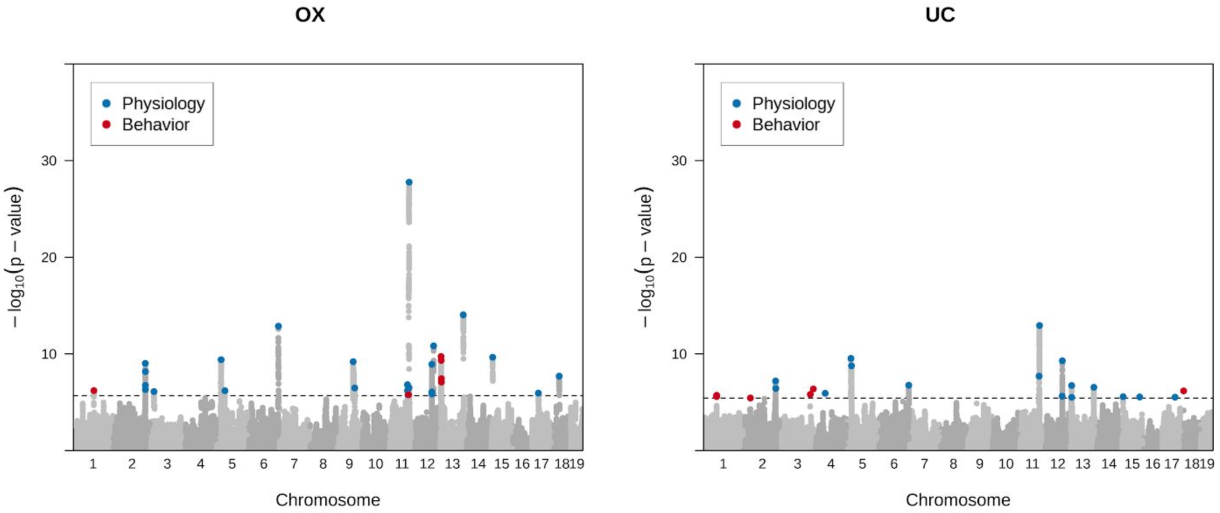

*Supplemental Figure 9: We estimated power of each QTL after correcting for Winner's Curse in the OX study, UC study, and mega-analysis ("OX", "UC", and "Mega", respectively). The distributions of the power levels in each study show that the mega-analysis obtained higher power than the component studies.*

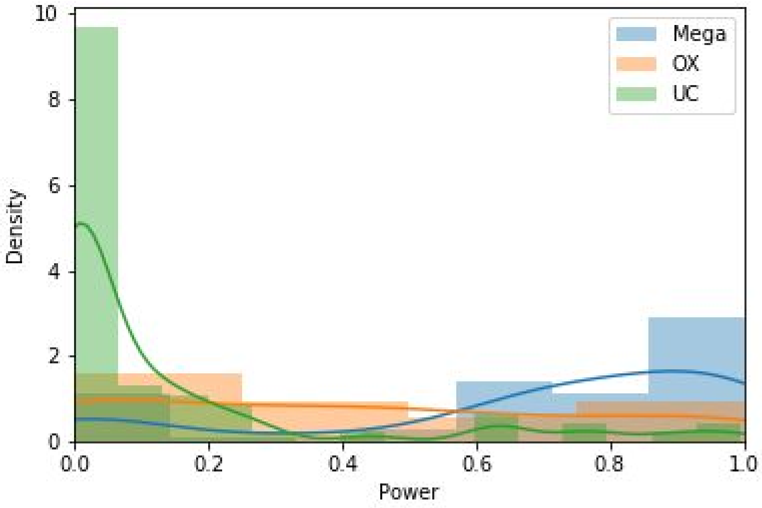

### Supplemental Tables

*Supplemental Table 1: Phenotype descriptions. We categorized the 23 phenotypes into several categories ("Category"). The sample sizes of the component studies are shown in the "OX" and "UC" columns. The phenotype abbreviations ("id") are used in Supplemental Data Files.*

| Phenotype name | Category | Description | OX | UC | id |
|---|---|---|---|---|---|
| startle habituation ratio | prepulse inhibition | Habituation (calculated as ratio of reduction from first to last startle stimulus) | 1682 | 824 | habit.ratio |
| startle habituation difference | prepulse inhibition | Habituation (calculated as difference first to last startle stimulus | 1703 | 983 | habit.diff |
| startle | prepulse inhibition | Startle | 1706 | 962 | startle |
| PPI with +6db prepulse | prepulse inhibition | PPI with +6db prepulse | 1720 | 936 | pp6.ppi |
| PPI with +12db prepulse | prepulse inhibition | PPI with +12db prepulse | 1725 | 946 | pp12.ppi |
| corrected freezing to context | fear conditioning | % time freezing during context test (corrected for baseline) | 1866 | 1025 | fc.context.corr |
| corrected freezing to tone alone | fear conditioning | % time freezing to tone prior to shock (corrected for baseline) | 1873 | 1035 | fc.uncond.freeze.corr |
| corrected freezing to cue | fear conditioning | % time freezing during cue test (corrected for baseline) | 1904 | 1030 | fc.cue.corr |
| baseline freezing D1 | fear conditioning | % time freezing at baseline during training | 1874 | 1035 | fc.baseline |
| activity decay | locomotor activity | Decrease in activity from beginning to end | 1899 | 954 | decay.activity |
| locomotor activity initial | locomotor activity | Activity from 0-15 min | 1902 | 954 | activity.begin |
| total locomotor activity | locomotor activity | Total activity from 0-30 minutes | 1900 | 954 | activity.total |
| locomotor activity end | locomotor activity | Activity from 15-30 minutes | 1900 | 954 | activity.end |
| tail length | physiology | Tail length (cm) | 1549 | 1053 | tail.length |
| TA | physiology | Weight of tibialis anterior (mg) | 1828 | 1057 | TA |
| soleus | physiology | Weight of soleus (mg) | 1832 | 1056 | soleus |
| gastroc | physiology | Weight of gastrocnemius (mg) | 1832 | 1054 | gastroc |
| plantaris | physiology | Weight of plantaris (mg) | 1832 | 1056 | plantaris |
| EDL | physiology | Weight of extensor digitorum longus (mg) | 1833 | 1054 | EDL |
| BMD | physiology | Bone mineral density | 1838 | 960 | bmd |
| tibia | physiology | Length of tibia (mm) | 1841 | 1022 | tibia |
| Abnormal BMD | physiology | Dichotomized bone mineral density | 1859 | 960 | bmd.a |
| weight | physiology | Weight (g) | 1865 | 1038 | weight |

*Supplemental Table 2: Heritability of phenotypes mapped in the mega-analysis. Heritabilities ("heritability") were determined using pruned genetic variants with INFO score > 90. The standard errors of the heritability estimates are shown in column "standard error".*

| phenotype | h2 | h2_se |
|---|---|---|
| corrected freezing to cue | 0.05 | 0.02 |
| PPI with +12db prepulse | 0.05 | 0.02 |
| corrected freezing to context | 0.06 | 0.02 |
| Abnormal BMD | 0.06 | 0.02 |
| startle | 0.08 | 0.02 |
| gastroc | 0.15 | 0.02 |
| locomotor activity end | 0.15 | 0.02 |
| tail length | 0.15 | 0.02 |
| locomotor activity initial | 0.17 | 0.02 |
| BMD | 0.18 | 0.02 |
| total locomotor activity | 0.18 | 0.02 |
| plantaris | 0.18 | 0.02 |
| weight | 0.19 | 0.02 |
| soleus | 0.19 | 0.02 |
| EDL | 0.21 | 0.02 |
| TA | 0.22 | 0.02 |
| tibia | 0.33 | 0.03 |

Supplemental Table 3: 95% confidence intervals of QTLs identified in mega-analysis. For each QTL in the mega-analysis, we compiled the estimated effect size in the mega-analysis ("beta"), the standard error on the effect size ("se"), and the negative logarithm of the p-value ("logP"). We also computed 95% confidence intervals on each QTL. The starting position of the confidence intervals are in the "from.bp" column, and the ending position of the confidence intervals are in the "to.bp" column. The coordinates are 1-indexed and inclusive.

Supplemental Table 4: Nonsynonymous mutations identified through fine-mapping of mega-analysis QTLs. We identified variants implicated by the SusieR fine-mapping method that were nonsynonymous mutations in genes. These genes are likely involved in the tested traits and in many cases have been independently identified in similar GWAS studies or mouse knockout studies ("GWAS Phenotype" and "KO Mouse Phenotype"). The references for these previously published studies are provided in the "References" column. The lead QTLs are shown in the "QTLs" column, and the confidence levels for the QTLs are shown in the "confidence column". 1 corresponds to the highest level of confidence, and 4 corresponds to the lowest level of confidence. The confidence levels were obtained using a combination of replication data, estimated level of confounding, and power in the mega-analysis.

Supplemental Table 5: Colocalization of mega-analysis QTLs. For each QTL in the mega-analysis, we computed the effect size, standard error on the effect size, and log p-value for every other trait (columns "beta", "se", and "logP", respectively). The phenotype in which the QTL was found in the mega-analysis is in the "phenotype_qtl" column, and the other phenotypes that colocalize are in the "phenotype" column. We used a Bonferroni corrected threshold of 0.05 to identify QTLs that significantly colocalized with multiple traits.

Supplemental Table 6: Confidence levels of QTLs identified in mega-analysis. For each QTL in the mega-analysis, we compiled the estimated effect size in the mega-analysis ("combined.beta") and the standard error ("combined.se"). We also computed the negative logarithm of the p-values for the mega-analysis and component studies ("combined.logp", "ox.logp", "uc.logp"). We corrected the bias due to Winner's Curse and used the corrected effect size estimates to compute the power of the association in each study ("combined.power", "ox.power", "uc.power").

### Supplemental Data

*Supplemental Data 1: Phenotypes*

*Supplemental Data 2: Genotypes*

*Supplemental Data 3: Genome-wide Association Studies*

*Supplemental Data 4: Fine-mapping (lists of SNPs in causal set for each QTL)*
